# Synovial fluid glycosaminoglycans signature involved 3-O-sulfated heparan sulfate modulated by TGF-β pathway in OA

**DOI:** 10.64898/2026.10.22.760925

**Authors:** Benjamin Even, Chayma Saadan, Camille Flageollet, Amina Boukhobza, Tarik Ould-Braham, Sara Shamdani, Etienne Gazaix-Fontaine, Isabelle Sacco, Sophie Cheverry, Stephanie Teboul, Xavier Chevalier, Florent Eymard, Patricia Albanese

## Abstract

Osteoarthritis (OA) involved extracellular matrix (ECM) remodeling of cartilage associated to inflammatory activation of fibroblastic like synoviocytes (FLS) from the synovium, but the molecular crosstalk between these two compartments remains poorly understood. Among Glycosaminoglycans (GAG) compound from ECM, heparan sulfate (HS) specifically regulate the bioavailability and activity of Heparin Binding Proteins (HBP), including cytokines and growth factors, such as TGF-β, whose aberrant bioavailability in the synovial fluid (SF) could drive the catabolic process. Here, we investigated the composition, sulfation patterns and biological activities of GAG from OA SF that are currently uncharacterized and explored the regulation of 3-*O-*sulfated HS in TGF-β mediated response in FLS.

Total GAG from 52 OA-SF SF were extracted and quantitatively characterized by DMMB assay and HPLC. They are functional since they are able to bind several HBP with pleiotropic inflammatory and angiogenic properties according to ELISA competitive binding assay. They were able also to transactivate HBP signaling in cellular tests in-vitro, since they potentiated proliferative properties on fibroblastic and endothelial cells, and induced altered metabolic and inflammatory markers expressions in murine chondrocytes and human FLS. Although HS represented a minor fraction of SF-GAG, they contained a rare tri-sulfated disaccharidic fraction distinct from GAG from cartilage. Same GAG characterization was performed on FLS expanded from SF and SM samples. FLS exhibited also a rare sulfation pattern of 3-*O-* Sulfated HS on their surface analyzed by immunofluorescent staining, associated with the expression of 4 isoforms of 3-*O*-sulfotransferase (HS3ST) identified by RT-qPCR. TGF-β1 treatment of FLS culture increases HS3ST expression profiles and HS sulfation pattern through the SMAD2/3 pathway, as observed by RT-qPCR, IF and western blot respectively. Finally, combined siRNA mediated silencing of these HS3ST enzymes affects FLS inflammatory secretome, according to proteome profiler cytokine arrays. among pro-inflammatory CXCL10 and CCL5 chemokines.

Altogether, these data identify OA SF-GAG as new effective modulator of joint cell responses and reveal for the first time 3-*O*-sulfated HS and HS3ST patterns of expression in OA FLS associated with TGF-β-Smad2/3-signaling pathway. Such specific HS fractions may contribute to the regulation of synovial inflammatory secretome, according to HBP partners that need to be further identified. However, this new glycanic mechanistic pathway sustaines the involvement of HS in OA progression and subsequently their interest as glycanic therapeutic targets.

## INTRODUCTION

Osteoarthritis (OA) is characterized by degradation of extracellular matrix (ECM) components of articular cartilage, considered for a long time as a central event. However, recent studies indicate that the synovial membrane (SM) also plays an important role in OA (Sellam et Berenbaum 2010; Shen et al. 2024; Zhao et al. 2023). It is composed of two main residents cell types: macrophage-like synoviocytes (MLS) which clear cartilage debris and participate in immune responses and fibroblast-like synoviocytes (FLS) producing synovial Fluid (SF) and the ECM of the synovial intima, including proteoglycans (PG) carrying heparan sulfate (HS) chains like perlecan and syndecan (Berenbaum 2013). SM changes and SF effusion in joint are correlated with inflammation and degree of OA and occur early in many patients (Sarmanova et al. 2016). The temporal sequence of pathological events within the joint during the OA process remains unclear. It is not known how early SM alterations shape the onset and progression of OA, nor how the synovium-cartilage crosstalk contributes to disease mechanism. Notably, pro-inflammatory cytokines and growth factors produced by FLS can induce a catabolic phenotype in chondrocytes (Fernandes et al. 2002). Among them, TGF-β plays an ambivalent role: it is crucial for maintaining healthy cartilage homeostasis but can contribute to pathological changes in aged or inflamed cartilage, according to concentration and biodisponibility of its active form (van der Kraan 2017; Cherifi et al. 2021). In a normal synovial joint, the chondrocyte’s pericellular matrix sequestered latent form of TGF-β through interactions with Perlecan HS-PG and Betaglycan (TGFbR3) whereas active TGF-β is only present after joint loading for a short period (X. Tang et al. 2018). In contrast, permanent and high levels of proteolytically active TGF-β are detected in SF from OA joints (Van Der Kraan 2018). This suggests that ECM and SF components such as PG and their associated glycosaminoglycans (GAG) chains could take part in synovium-cartilage crosstalk through crucial control of Heparin Binding protein (HBP) levels such as active TGF-β.

GAG are linear and sulphated polysaccharides divided in distinct families, heparan sulfate (HS), chondroitin sulfate (CS), dermatan sulfate (DS), or keratan sulfate (KS), according to the (Glc and/or Gal) composition of their very long chains based on repeated disaccharidic units of uronic acid and hexosamine. Among them, CS and un-sulfated Hyaluronic Acid (HA) are major ECM components of cartilage and SF and their involvement in healthy mechanical properties and turnover during OA are described for a long time. Thus, degradation of Aggrecans during OA (Huang et Wu 2008), the main cartilage PG associated with numerous CS, KS and HA chains, have paved the way of therapeutical strategies based on CS and HA visco-supplementation. However, even if present in low amounts, we and others recently associated HS from cartilage to important effects during OA pathophysiology: rare cartilaginous HS evolved in amount, structure and functionality during OA and are able to modulate chondrocyte phenotypes according to specific modification of their 6-*O*-sulfation pattern (Chanalaris et al. 2019; Severmann et al. 2020; Shamdani et al. 2020).

Indeed, HS have a higher structural diversity among GAG families, associated to functional complexity. HS disaccharide backbone is composed of 6-carbon sugars that are Glucosamine (GlcN) linked to Glucuronic acid (GlcA) or its epimerized Iduronic acid (IdoA) form. HS chains are further characterized by alternating domains with different levels of *N*- or *O*-sulfation (SO) substituted respectively on the carbon 2 (C2)’s amino group (-NH) of GlcN, and on the hydroxyl groups (-OH) of GlcA/IdoA’s C2 and GlcN’s C6 and C3. This complex HS sulfation profile is driven by post polymerisation modification in Golgi apparatus due to different enzymes isoforms named sulfotransferases (HS-x-ST). These sulfation patterns defined the HS structural scaffold required for selective binding of heparin-binding proteins (HBP) such as growth factors or cytokines (Murphy et al. 2004; Alotaibi et al. 2024). Through these interactions, HS can sequester and compartmentalize HBP within ECM niches, thereby protecting them from proteolytic degradation and thereby regulating their diffusion, local bioavailability and cellular effects. Whereas 2-*O*- and 6-*O*-sulfated structures in HS chains are ubiquitous and involved in several signalling pathways such as those of FGFs (Chan et al. 2015) and WNTSs (Gao et al. 2016), 3-*O*-sulfated motifs are rarer and described as “non-essential”, as genetic disruption of HS3STs leads at most to a milder phenotypes. However, they were proposed to be associated with disease vulnerability and progression (Ouidja et al. 2024), such as Alzheimer’s disease (Sepulveda-Diaz et al. 2015), breast cancer (Hellec et al. 2018) and during viral infection (Shukla et al. 1999). However, little is known about their specific interactions with HBP ligands and their effects on signaling pathways.

As rare in joint tissues, few are known on HS fundamental roles in homeostasis, as well as in orchestrating pathological processes such as inflammation, angiogenesis, and tissue remodeling in OA (Ricard-Blum et al. 2024). Thus, we proposed to explore HS chains expressions in SF, SM and FLS from human OA samples, their sulfation patterns and associated sulfotransferase expressions, and HS functionality such as HBP binding and effects on joint cells properties, alone and associated with TGF-β signaling pathway.

## MATERIALS AND METHODS

### Human samples collection

***SF, SM and Blood samples from OA Patients*** were recruited from orthopedic and rheumatologic departments at Henri Mondor’s Hospital (Créteil, Val de Marne, France), within the framework of a clinical protocol approved by the Hospital’s Ethics Commission N° 07-34 and from the BIOGO cohort (ethic committee Sud Ouest et Outre Mer II.2022-A00685-38), after informed consent of the patients. SF were obtained by arthrocentesis from patients with painful knee OA and synovial tissues were isolated from patients undergoing total knee replacement surgery for symptomatic knee OA. Demographic and radiological characteristics were collected at enrollment. Radiographs were scored according to the Kellgren and Lawrence (KL) classification. ***Human cartilage samples from control and OA patients*** were obtained and analysed as previously described (Shamdani et al. 2020)**. *Blood samples*** from healthy donors (CT) were obtained through an agreement with “*Etablissement Français du Sang*” (EFS), following the Ethical approval n° 2022-2026-027 CCPSL UPEC.

### GAGs extraction and quantification from synovial fluid, serum and cartilage

GAG extraction was performed according to a modified version of (Huynh, Villares, et al. 2012) protocol described in supp. data. GAG extraction form cartilage were performed as described in (Shamdani et al. 2020). Total sulfated GAG, CS and HS families were quantified according to 1-9 dimethyl-methylene blue (DMMB, Sigma Aldrich) assay and after specific enzymatic digestions (chondroitinase) and chemical treatment (nitrous acid) of GAG respectively, according to (Barbosa et al. 2003) protocol reported in supp data. GAG were used further for HPLC analysis, Elisa competitive assay on HBP and functional tests on cells described in (Huynh, Villares, et al. 2012; Shamdani et al. 2020) and reported below or in supp data respectively.

### Isolation of Fibroblast-like synoviocytes (FLS) from SF and SM, cultures conditions and treatments

FLS were isolated from SF and SM according to an adapted protocol based on (Scanu et al. 2007) and (Rosengren et al. 2007). Briefly, SF were treated with hyaluronidase (10 UI/mL, Sigma-Aldrich, 45 min, 37°C) then diluted in PBS (pH 7-7.2, with calcium and magnesium, FisherScientific) with calcium and magnesium and centrifuged (300 g, 10 min). Concerning SM, tissue samples were digested in complete medium (DMEM low glucose, GlutaMAX™ Gibco, 10% (v/v) FBS (Cytiva, Hyclone); 1% Penicillin/Streptomycin (PS 10,000 U/mL, FisherScientific) with collagenase type VI (1 mg/mL Sigma-Aldrich, 12 hours, 37 °C, 5% (v/v) CO), and dissociated cells were centrifuged at 250 g, 10 min. Both SF and SM-FLS pellet were resuspended in complete medium, seeded at high density (1 x 10^6^ cells/cm^2^) and cultured until confluent clones appeared. Cells were sub-cultured with trypsin–EDTA and used between passages 0–7. All treatments were performed on FLS between passages 2 to 4, preliminary seeded at 2,500 cells/cm², grown to at least 80% confluence and left for 24 h in serum-starved medium (DMEM, 1% PS). Treatments were further performed with recombinant human TGF-β1, IL-1β, FGF2, FGF18 (commercial references in Supp. Table S1) or pre-filtered GAG extracted from OA-SF, as described in the figure legends. To analyze the TGF-β pathway, a specific inhibitor of Smad2/3 (SB-431542, SIGMA) or Smad1/5/9 pathway (ML-347, Sigma) were included. After PBS washes, cells were incubated for 24 h in serum-free depletion medium (DMEM, 1% PS, 1 mg/mL BSA), and these conditioned media were collected, centrifuged (4,696 g, 10 min) and stored at −80 °C for ELISA assays.

### Immunofluorescence analysis of FLS *in vitro*

SF- and SM-FLS were seeded at 10 000 cells/well in labtek slides, fixed 10min in 4% (w/v) paraformaldehyde (PFA, Thermo Scientific Alfa Aesar), and washed 3 times with PBS. When intracellular staining was required, cells were permeabilized for 10 min with 0.1% (v/v) Triton X-100 in PBS. After blocking 1h (RT) in PBS with 3% (w/v) BSA (Sigma-Aldrich) and 100 mM glycine, primary antibodies were applied in PBS with 3% (w/v) BSA overnight (Supp. Table S3). Conventional primary antibodies were detected with donkey anti-rabbit Alexa Fluor-488 (1:250, Invitrogen) for 1 h at room temperature. When single-chain ‘phage display (tagged) antibodies were used (Lensen et al. 2005), an anti-VSV (rabbit) step (1:5,000) was performed for 1 h at room temperature, followed by the donkey anti-rabbit Alexa Fluor-488 secondary (1h). Slides were counterstained with DAPI (Invitrogen), mounted in ProLong Gold Antifade and imaged on an inverted microscope with disk-scanning confocal unit (IX81/IX2-DSU, Olympus). For each experiment, 6–8 fields/images per condition were acquired consecutively using identical acquisition settings. Fluorescent signal was quantified further using Fiji/ImageJ (NIH, USA): For each image, the total fluorescence signal of the entire field of view was measured as the Integrated Density. Background fluorescence was determined from a cell-free region of the same image and used to calculate the Corrected Total Cell Fluorescence (CTCF = Integrated Density− (Image Area×Mean Background Fluorescence)). The CTCF value obtained for each field was then normalized to the total number of cells present in the corresponding field: CTCF per cell = CTCF / Total number of cells.

### Statistical analysis

Patient characteristics are described using frequencies (%) for categorical variables and mean ± SD or median (Q1-Q3) for continuous variables. Experimental data are presented as mean ± standard error of the mean (SEM). Data distribution was assessed using the Shapiro-Wilk test, prior to statistical analysis. Comparisons between two independent groups were performed using the Mann-Whitney U test. Paired comparisons were performed using either paired Student’s t-tests or Wilcoxon signed-rank tests, according to normality test. Comparisons among multiple groups were performed using one-way analysis of variance (ANOVA) followed by Dunnett’s or Bonferroni’s post hoc tests when data met the assumptions for parametric analyses. For non-parametric data, comparisons were performed using the Kruskal-Wallis test followed by Dunn’s multiple-comparison test. Correlations between continuous variables were assessed using Spearman’s rank correlation coefficient.

All analyses were conducted using GraphPad Prism 8 software. Statistical significance was defined as p < 0.05 and indicated as follows: p < 0.05 (*), p < 0.01 (**) and p < 0.001 (***).

## RESULTS

### Glycanic characterization of synovial fluid

A total of 52 OA synovial fluid (SF) samples were included in this study for GAG extraction and further FLS cultures (clinical data described in Fig. S1A). Sulfated GAG concentrations from SF were comprised between 10 and 60µg/ml suggesting an important inter donor variability, and almost 2-fold higher than that found in healthy and OA patient serum. There was no difference in SF-GAG concentrations according to gender (Fig. 1C), or any correlation with age (Fig. 1D), Kellgren and Lawrence (KL) score (Fig. 1E), and Lequesne score (Fig. 1F). No significant Spearman correlations were observed between total SF-GAG and several HPB concentrations, such as TGF-β, FGF2 or Il-1 β, that were measured as OA mediators (Boffa et al. 2021; De Roover et al. 2023) in the same OA SF (Fig S1, B-C).

**FIGURE 1:**
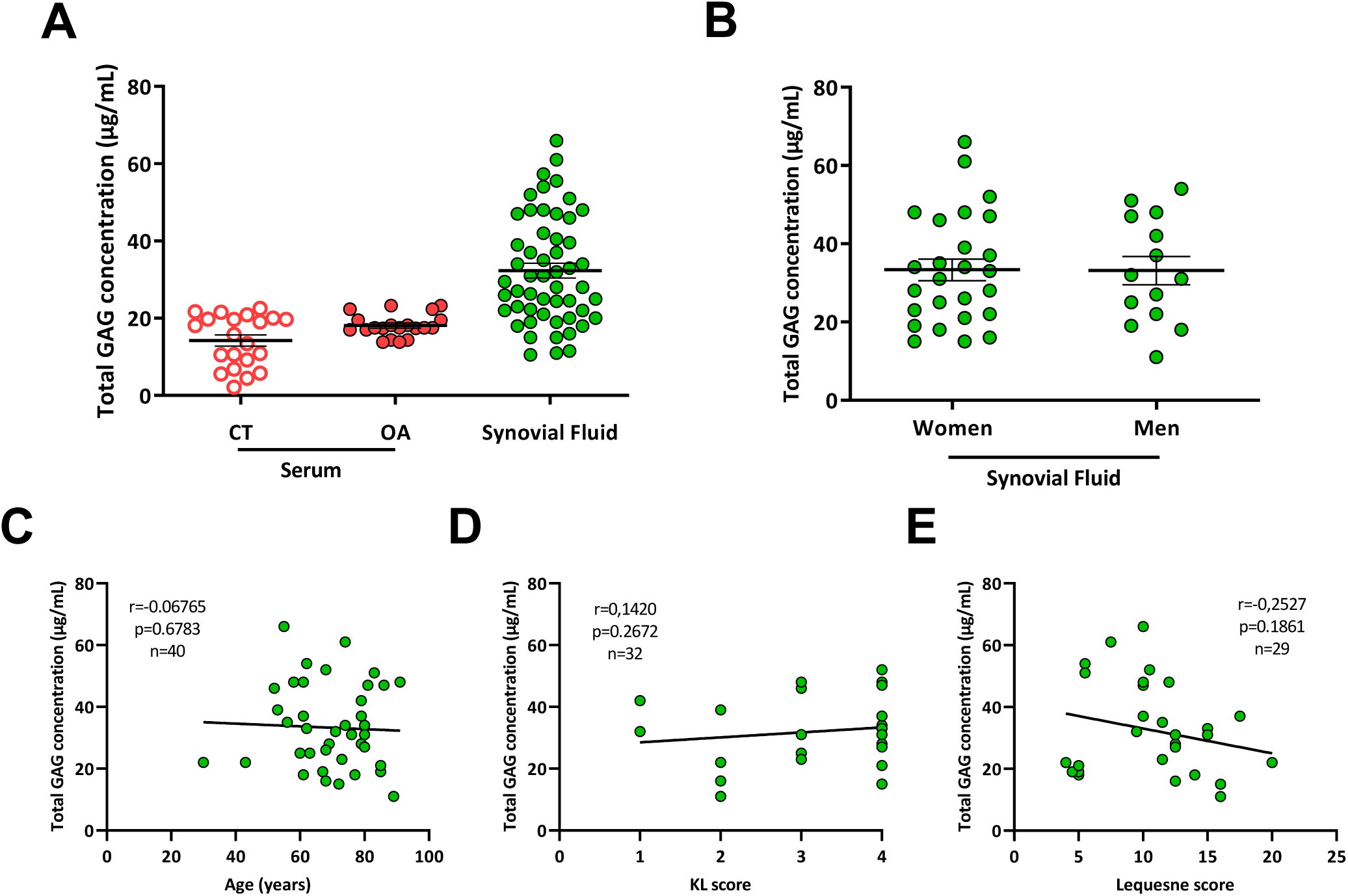
Glycanic characterization of Synovial Fluids. (A) Total sulfated GAG were extracted from biological fluid (healthy control serum (CT) (n = 21), OA Serum (n = 10) and OA synovial fluid (n= 40) and quantified by the DMMB assay. Results are expressed as µg of GAG per mL of biological fluids. (B) Comparison of total sulfated GAG concentration in SF (SF-GAG) by sex (Mann-Whitney statistical t test). Relationship between SF-GAG concentration and (C) donor age; (D) Kellgren-Lawrence (KL) score and (E) Lequesne index. Spearman’s rank correlation, p-value and n reported.

SF-GAG showed very high binding affinity to PDGF-bb and TGF-β exceeding 8 and 2 times those of heparin, respectively (Fig 2A). By contrast, binding affinity to PTN, VEGF, Il-6 and FGF-2 were lower than heparin and heterogeneous across SF samples. We then tested whether these GAG-HBP interactions were translated into receptor activation through distinct cellular tests. VEGF-induced proliferation of HUVEC was potentiated by some SF-GAG leading to a significant increase in the mitotic index, exceeding mean value observed with heparin (> 152%), whereas other SF-GAG had no additive effect on VEGF-induced proliferation (mitotic index < 152%) (Fig. 2B). The FGF-2 mitotic index in BAF32 cells was also increased in presence of heparin and some SF-GAG (mitotic index > 173%) (Fig. 2C) whereas other SF-GAG had little or no effect. SF-GAG were tested alone on primary mice chondrocytes (Fig. 2E-F): some SF-GAG upregulated gene expression of MMP13 catabolic marker and downregulated ACAN anabolic marker compared to unstimulated cells, with changes more pronounced than those induced by IL-1β, whereas some other induced minimal change. These suggested that SF-GAG are functional since some of them interact with HBP and/or potentiate HBP effects on different cells from the joint, with inter-donor differences.

**FIGURE 2:**
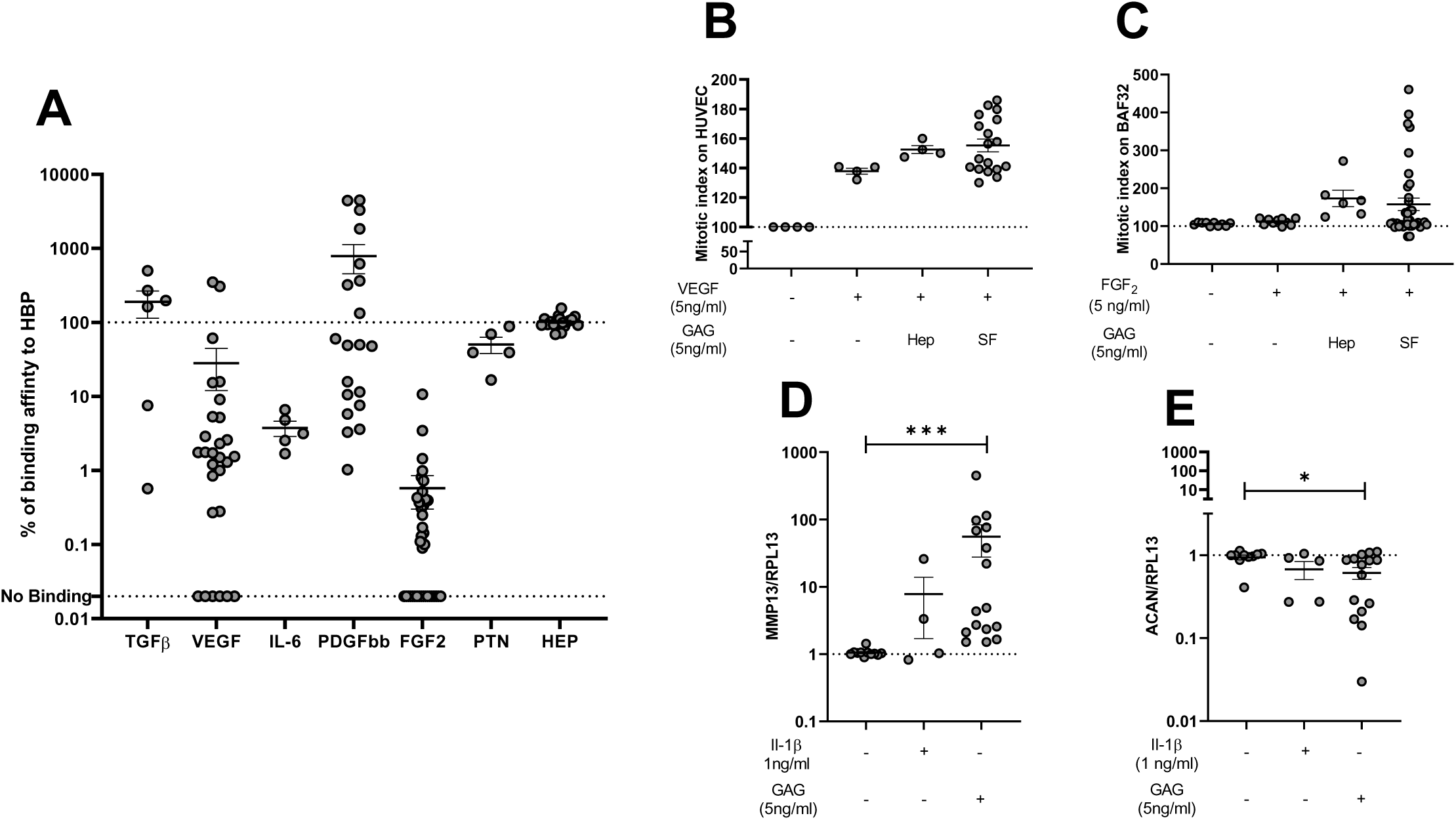
Functional properties of GAG extracted from OA synovial fluid. (A) Percentage (%) of binding affinity of SF-GAG to selected heparin-binding proteins (HBP). SF-GAG binding affinity is expressed as a percentage relative to heparin (HEP), which was set to 100% of binding affinity (see Methods). Each point is an individual SF sample (results shown as mean +/- SEM). (B) Mitotic index of HUVEC after 48 h in the presence of with VEGF (5 ng·mL⁻¹) with or without SF-GAG (5ng·mL⁻ ¹). Values are normalized to control cells (no VEGF, no GAG = 100%). Proliferation was assessed using PrestoBlue™ reagent. (C) Mitotic index of BAF32 cells after 48 h in the presence of FGF2 (5 ng·mL⁻¹) with or without SF-GAG (5 ng·mL⁻¹). Values are normalized to control cells (no FGF2, no GAG = 100%). BAF32 cells require heparinoid cofactors to respond to FGF2 (see Methods). Proliferation was assessed using MMT assay. (D) and (E) RT-qPCR quantification of catabolic (MMP13) and anabolic (ACAN) marker expression respectively in primary murine chondrocytes treated with SF-GAG (5 ng·mL⁻¹) or IL-1β (1 ng·mL⁻¹) for 24 hours. Expression is shown as fold change versus untreated control (mean ± SEM). Statistical tests: Mann-Whitney statistical t test, *p < 0.05, ***p < 0.001.

### Functional characterization of SF GAG on synoviocytes properties

Next, we tested the impact of SF-GAG on FLS proliferation induced by different inflammatory mediators (TGF-β, IL-1β, FGF-2, and FGF18) (Fig. 3). Sub-optimal mediator concentrations that stimulated FLS proliferation were first identified for TGF-β (20 ng/ml), Il-1β (0.1 ng/ml), FGF2 (5 ng/ml) and FGF18 (10 ng/ml) (Fig 3A). Then SF-GAG (from 2 to 5 donors of SF samples) were tested on 4 distinct FLS cultures, to measure the variability of the SF-GAG response with respect to the FLS culture donors (4 distinct colors) (Fig. 3B-F). These data indicated that low concentrations of SF-GAG (0.01 to 0.1 µg/ml) significantly reduced the proliferation of FLS (Fig. 3B). Moreover, these SF-GAG concentrations significantly attenuated FLS proliferation induced by TGF-β, Il-1β, FGF-2, and FGF18 (Fig. 3C-F). In parallel, inflammatory responses of FLS were assessed (Fig. 4) and indicated that IL-1β and TGF-β strongly increased Il-6 and Il-8 secreted levels whereas FGF18 produced no consistent effect. Interestingly, SF-GAG alone reduced basal IL-8 secretion and had no impact on IL-6 secretion (Fig. 4C– D). In co-treatment experiments, SF-GAG did not consistently modulate Il-1β-induced secretion, but significantly increased TGF-β-induced secretion of Il-6 and Il-8 (Fig. 4C–D). These data indicated that SF-GAG can modulate inflammatory output of growth factors such as proliferative response and cytokines secretion by FLS.

**FIGURE 3:**
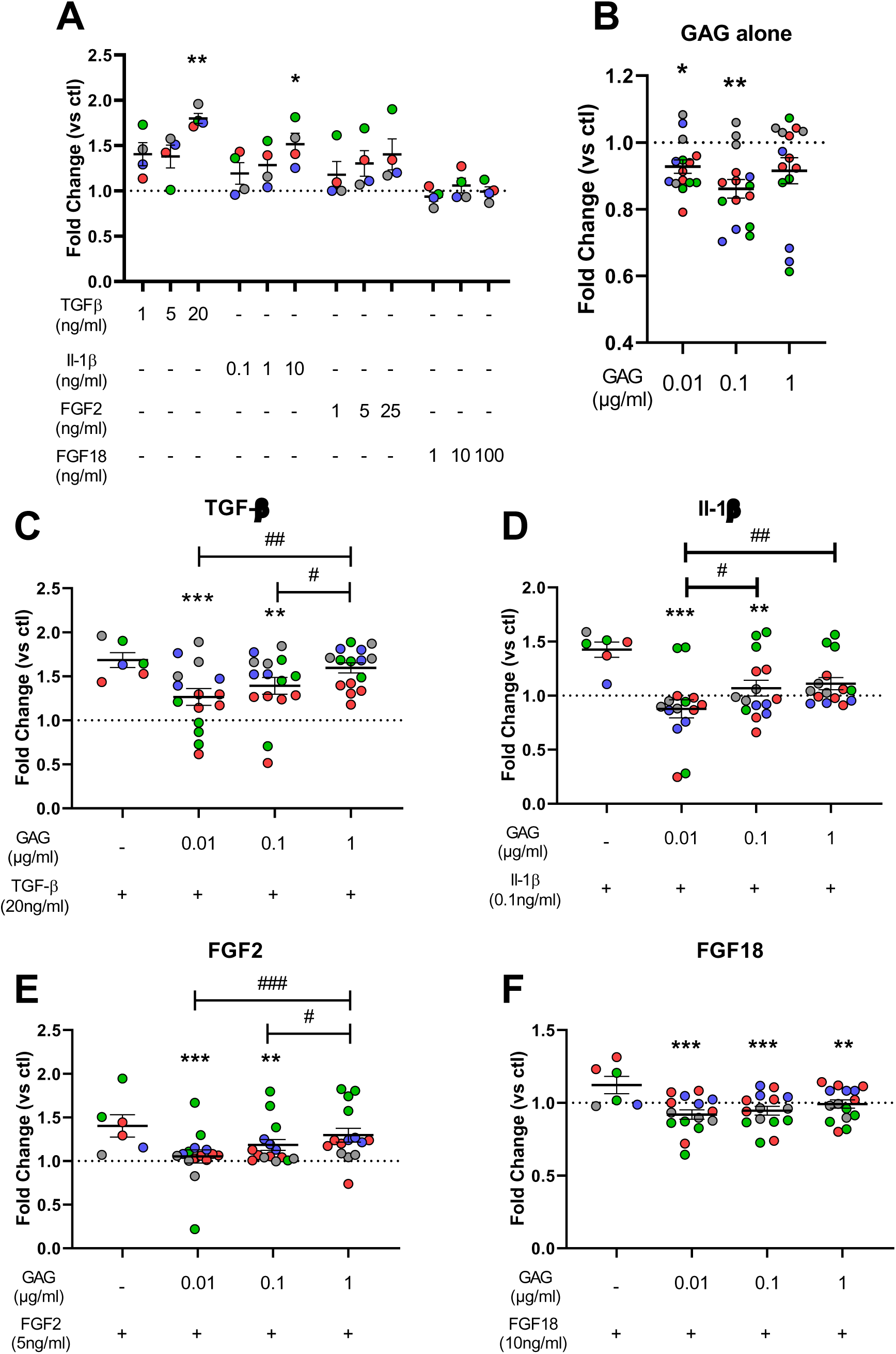
Effect of GAG from OA SF on FLS proliferation capacity induced by growth factors. (A) Dose–response of OA FLS to growth factors: TGF-β (1, 5, 20 ng·mL⁻¹), IL-1β (0.1, 1, 10 ng·mL⁻ ¹), FGF2 (1, 5, 25 ng·mL⁻¹) and FGF18 (1, 10, 100 ng·mL⁻¹). (B) Dose–response of OA FLS to SF-GAGs alone (0.01, 0.1, 1 µg·mL⁻ ¹). (C–F) Co-treatment of OA FLS with SF-GAGs (0.01, 0.1, 1 µg·mL⁻¹) and the indicated growth factors: TGF-β (20 ng·mL⁻¹), IL-1β (0.1 ng·mL⁻¹), FGF2 (5 ng·mL⁻¹) and FGF18 (10 ng·mL⁻¹). Proliferation was measured after 48 h using PrestoBlue™ cell viability reagent and values are normalized to the control condition (CTL: cells without growth factor and without GAG; set to 100%). Data are shown as mean ± SEM. Statistical comparisons: *p < 0.05; **p < 0.01 versus corresponding growth-factor alone (One way Anova and Dunnett’s multiple comparisons test). #p < 0.05; ##p < 0.01 indicate comparisons specified in the panels (One way Anova and Bonferroni post-test). Each symbol color corresponds to FLS derived from a distinct patient (n=4); multiple SF-GAG samples from different donors (2-5 donors) were tested on each FLS line (see Methods).

**FIGURE 4:**
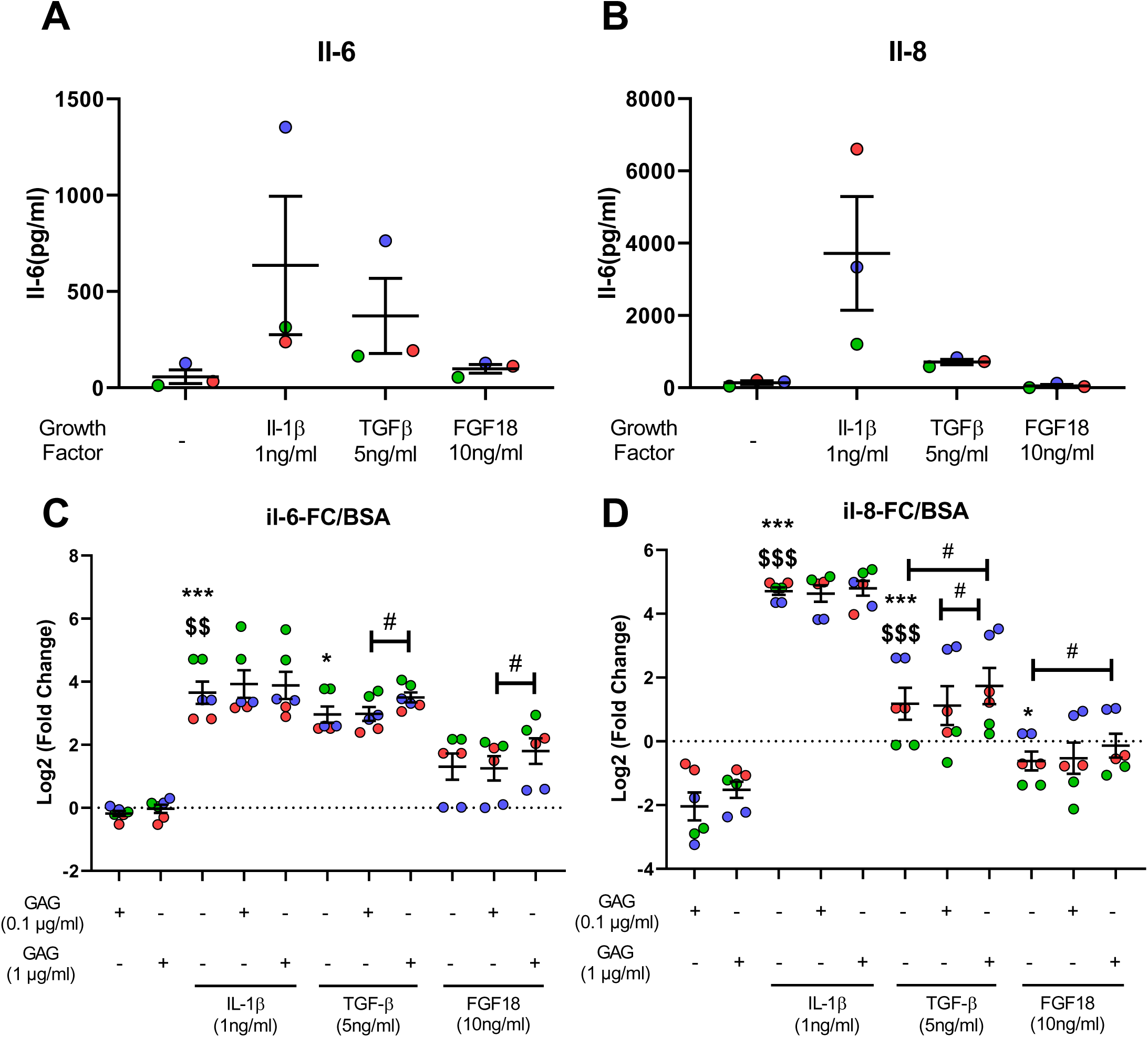
Effect of GAG from OA SF on FLS cytokines secretion induced by inflammatory mediators. (A–B) IL-6 (A) and IL-8 (B) concentrations in FLS culture supernatants measured by ELISA after 48 h of stimulation with IL-1β (1 ng·mL⁻¹), TGF-β (5 ng·mL⁻ ¹) or FGF18 (10 ng·mL⁻¹), (C–D). IL-6 (C) and IL-8 (D) concentrations in FLS culture supernatants measured by ELISA after 48 h stimulation with IL-1β (1 ng·mL⁻¹), TGF-β (5 ng·mL⁻ ¹) or FGF18 (10 ng·mL⁻¹) in the absence or presence of SF-GAG (1 and 0.1 µg·mL⁻¹) expressed as log2 fold-change relative to the untreated control (Log2[FC] = log2 (value / mean_control)). Data are presented as individual values with mean ± SEM. Each symbol color corresponds to FLS derived from a distinct patient (n=3); multiple SF-GAG samples from different donors were tested on each FLS. Statistical comparisons: *p < 0.05; ***p < 0.001 versus GAG (10-7 g/ml), $$p < 0.01 GAG (0,1 µg/ml) versus GAG (1 µg/ml), # p < 0.05 (One way Anova and Dunnett’s multiple comparisons test).

### Structural features of GAG species in OA

To provide a degree of structural understanding for the effects of the SF-GAG, the different GAG families present were analyzed as well as their sulfation profile. These data were compared to those obtained from GAG isolated from serum and cartilage of healthy and OA patients (Fig. 5). Details on the GAG family’s concentration (Fig 5A) indicate that the total GAG concentration significantly higher in SF as compared to serums (previously described in Fig. 1A), was due to higher concentrations in CS and other GAG species in SF as compared to serums. HS concentrations were low in OA SF and serum, to the same extend but with larger variability among SF. Fractions analysis (Fig. 5B) indicated that GAG from serum were mainly composed of CS (55%) and HS (45%), whereas GAG from SF are mainly composed of CS (68%), with lower amount of HS (16%) and other GAG species (KS and HA) (17%) These SF-GAG fractions were similar to those previously described in cartilage (Fig. 5B and (Shamdani et al. 2020)). GAG family members present different sulfation patterns (Fig 5C): un-sulfated for HA, sulfated on C2 of GlcA and/or (C4 and/or C6) of GalNac for CS, sulfated on C2 of GlcA/IdoA’s and/or C2, C6 and C3 of GlcN’s for HS. According to the heterogeneity of sulfation pattern available for HS disaccharides (Fig. 5D), we performed HPLC analysis to quantify the % of non-sulfated, mono, di- or tri-sulfated profile (Fig. 5E-F). CS and HS disaccharides from SF were mainly non-sulfated whereas CS and HS from cartilage (CT and OA) were mainly (mono- and di-) sulfated. Moreover, we identified a substantial level (5%) of tri-sulfated disaccharides of HS in OA-SF, that was not observed in other tissues. Thus, low amounts of HS from SF-GAG are characterized by very distinct sulfated profiles as compared to HS from cartilage. This specific tri-sulfated HS disaccharides fraction from SF-GAG suggests that these HS could have distinct tissular origin than cartilage and be of synovial origin.

**FIGURE 5:**
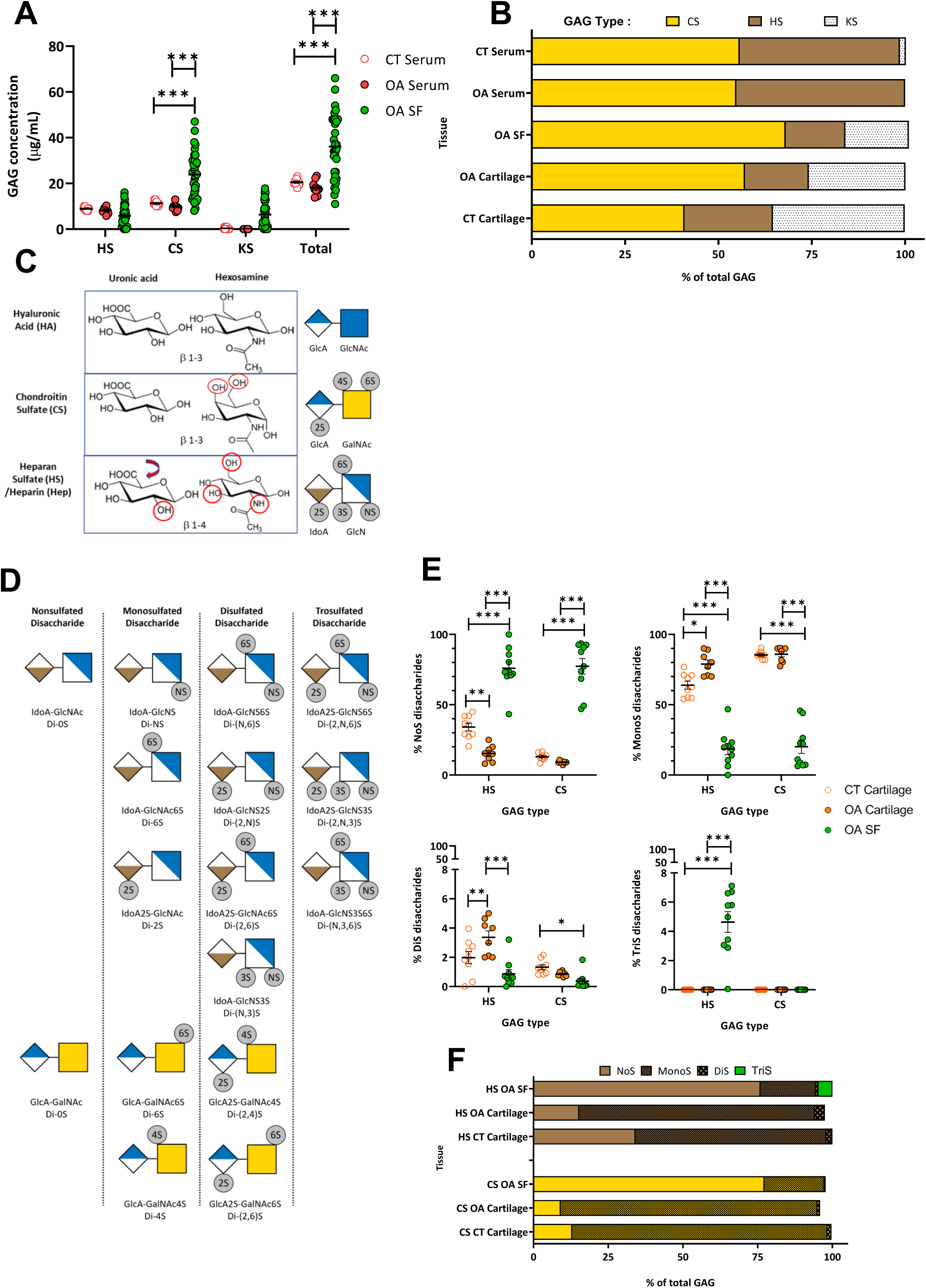
Structural characterization of HS, CS and others GAG species from healthy donors and OA Serum, Cartilage and Synovial Fluid. (A–B) Sulfated GAG family’s concentrations of HS, CS, and KS GAG, as well as to total, were measured in biological fluids: control serum (CT), OA serum (red), OA synovial fluid (SF, green) and expressed as µg.mL⁻ ¹. Values are mean ± SEM (Serum CT and OA: n = 10; SF: n = 32; Cartilage CT and OA: n = 9). (B) Histogram representation of relative contribution of each major GAG class (HS, CS and KS) to Total GAG (% of total GAG) in CT and OA serum, cartilage and SF. (C) Schematic representation of disaccharide units of Hyaluronic Acid (HA), Chondroitin Sulfate (CS) and Heparan Sulfate (HS) and associated possible sulfation positions (red circle). (D) Abbreviations and schematic key for disaccharide sulfation patterns used in panels E and F. (E) HPLC analysis of disaccharide sulfation pattern obtained after enzymatic digestion of HS and CS chains: non-sulfated (NoS), mono-sulfated (MonoS), di-sulfated (DiS) and tri-sulfated (TriS). Data are shown for cartilage (orange) CT (n = 9) and OA (n = 8) and SF (n = 10, green) as mean ± SEM.(F). Histogram representation of relative contribution of each of these 4 disaccharide sulfation patterns to HS and CS families (% of total GAG families). Statistical comparisons: *p < 0.05; **p < 0.01; *** pvalue<0,001 indicate comparisons specified in the panels (One way Anova and Bonferroni post-test).

### Evidence of a 3-O sulphated HS signature in FLS from OA patients

GAG were extracted from FLS cultured from SF and SM samples and were identified as an equal mix of HS and CS (47 and 53% of total GAG respectively, Fig. 6A and B). Distribution of pericellular matrix HS, analysed by immunofluorescence staining with antibodies recognizing different sulfated motifs (Fig. 6C), demonstrate that HS from SF-FLS and SM-FLS displayed a broad sulfation profile and were notably enriched in 3-*O*-sulfation (HS4C3 staining), consistent with the tri-sulfated disaccharides fraction analyzed by HPLC in SF-GAG (Fig. 4). These staining were decreased after FLS treated with heparinases confirming their selectivity for HS in these cells (Fig. S3 A). Finally, RT-qPCR analysis of the mRNAs encoding the isoforms of HS3STs involved in this sulfation pattern showed high levels of *HS3ST3A* and *HS3ST3B*, and low and heterogeneous levels *HS3ST1* and *HS3ST2* whereas *HS3ST5* and *HS3ST6* were undetected.

**FIGURE 6:**
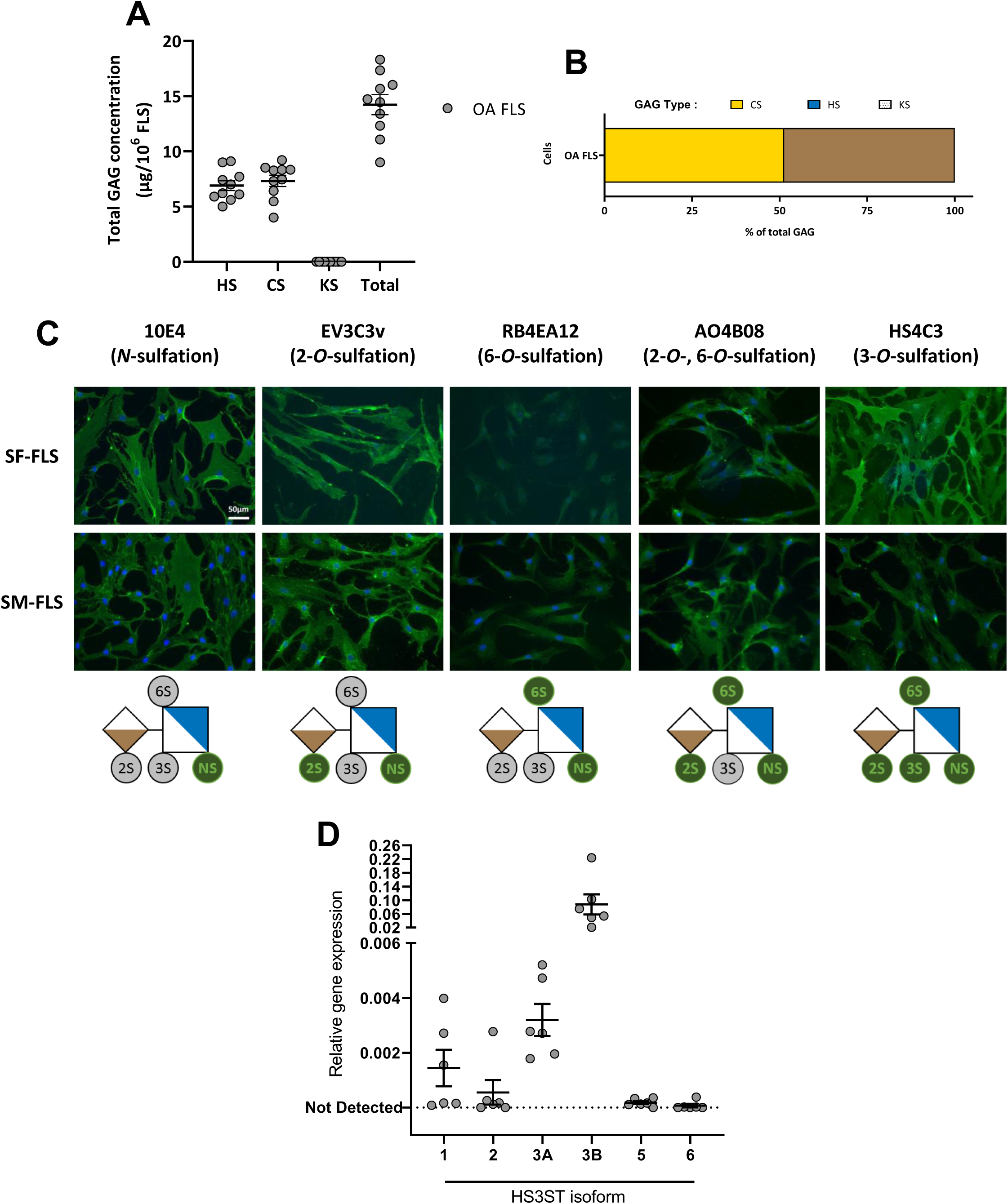
Structural characterization of GAG species in OA FLS. (A) Total sulfated GAG (HS, CS and KS) extracted from OA FLS, expressed as µg of GAG per 10^6^ cells. Data are represented as mean +/- SEM (n=10). (B) Relative proportion (%) of each GAG family (HS, CS, KS) in OA FLS (n = 10). (C) Representative immunofluorescence of distinct HS sulfation motifs labeled on OA FLS: 10E4 (N-sulfation), EV3C3v (2-O-sulfation), RB4EA12 (6-O-sulfation), AO4B08 (2- and 6-O-sulfation) and HS4C3 (3-O-sulfation). Nuclei were counterstained with DAPI. Scale bar = 50 µm. Lower panel, schematic key of HS disaccharide with sulfation patterns recognized by antibodies in green. HS4C3 recognizes highly sulfated HS epitopes containing 3-O-sulfated glucosamine, whereas AO4B08 recognizes epitopes associated with 2-O- and 6-O-sulfation. (D) Relative mRNA expression of HS3ST isoforms in OA FLS determined by RT-qPCR and expressed as 2^(-ΔCt) relative to α-tubulin (mean ± SEM, n = 6).

### TGF-β drives a robust increase in 3-O-sulfated HS on FLS through Smad 2/3 signalling pathway

As TGF-β1 was the most abundant HBP in OA SF (Fig. S1B) and had the strongest affinity to SF-GAG (Fig. 2A), we investigated if it could modulate the HS sulfation pattern in human OA FLS (Fig. 7). We first demonstrated that TGF-β1 selectively activated the canonical SMAD2/3 pathway in OA FLS cultures (Fig. S4, phosphorylation, and nuclear translocation). Subsequent immunofluorescent analysis (Fig. 7A-B) indicated that 2-*O* and 6-*O* sulfated HS present on these FLS were not modified with TGF-β1 induction. In contrast, the 3-*O*-sulfated motifs were increased on FLS cells treated by TGF-β1, and this labeling decreased when FLS were treated with SMAD2/3, but not with SMAD1/5/9, pathway inhibitor. Consistent with this observation, TGF-β1 induced a rapid and transient increase in *HS3ST1*, *HS3ST2*, *HS3ST3A,* and *HS3ST3B* mRNA expression, which was prevented by SMAD2/3 inhibition (Fig. 7C).

**FIGURE 7:**
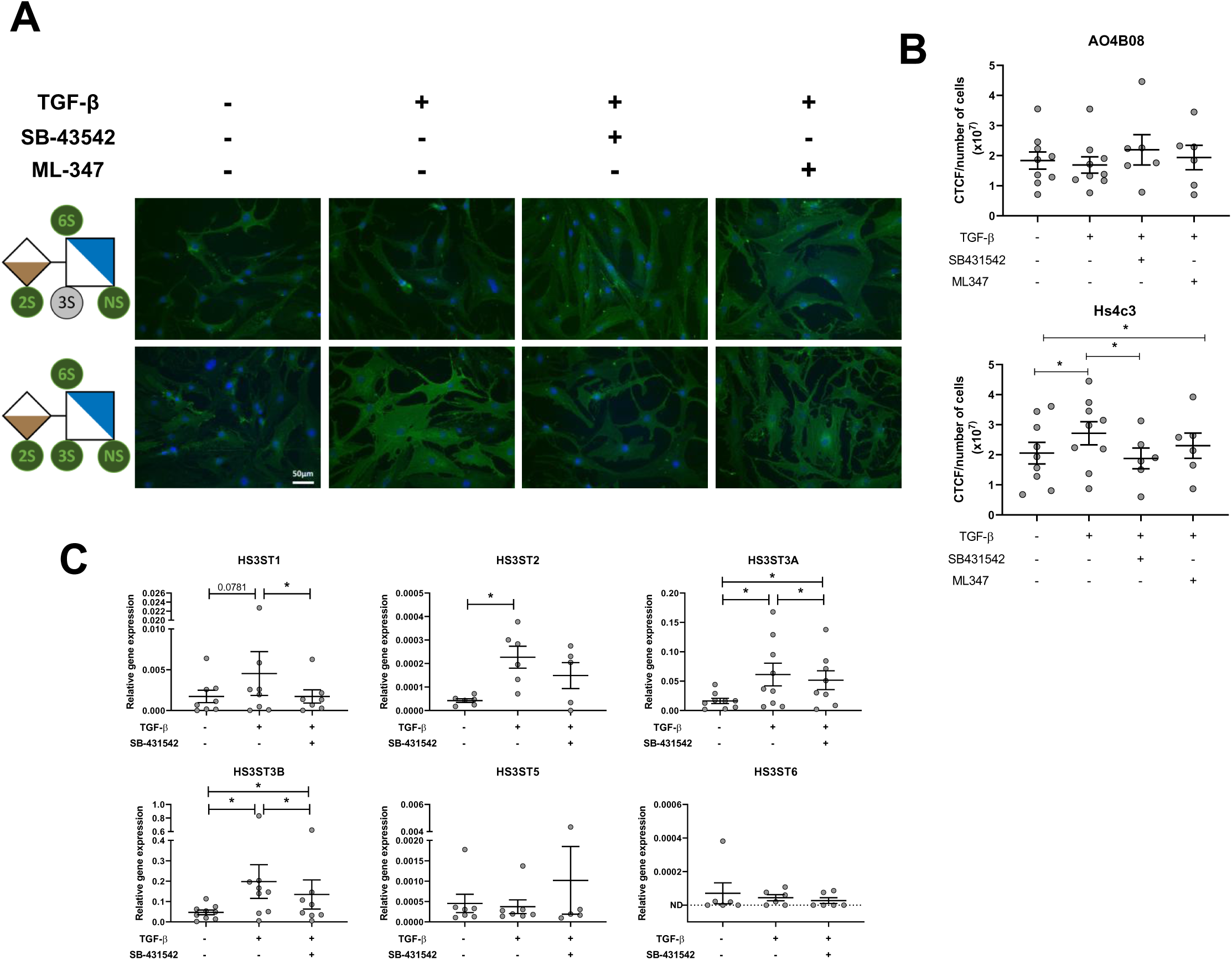
TGF-β modulates the 3-O-sulfation pattern via the Smad 2/3 pathway in FLS. OA FLS were treated with TGF-β1 (5 ng·mL⁻ ¹) ± SB431542 (10 µM) or ML-347 (1 µM). (A) Immunofluorescence (representative images) of HS4C3 (3-O-sulfation) and AO4B08 (2/6-O sulfation) staining (green) in 24h-treated FLS. Nuclei were counterstained with DAPI (blue). Scale bar = 50 µm. (B) Quantification of the corrected total cell fluorescence (CTCF) reported on the number of cells for HS4C3 and AO4BO8 staining (n=7). (C) RT-qPCR analysis of HS3ST isoforms expressions following 4h of treatment. Data are shown as relative expression (2^−ΔCt^, ΔCt = Ct_target − Ct_α-tubulin) and presented as mean ± SEM (n = 6–8). Statistical analysis: paired t-test was used for IF quantifications; Wilcoxon signed-rank test was used for gene expression data. p < 0.05; p < 0.01.

### siRNA knockdown of HS3ST reveals contribution of 3S-HS to FLS inflammatory response

To assess whether these HS could have an impact on the TGF-β effect on FLS inflammatory phenotype, we used a pooled of siRNA (siPool) to efficiently suppress the expression of all four of these HS3ST isoforms, and because individual HS3ST knockdown induced compensatory regulation of the remaining isoforms (Fig 8A). OA FLS were transfected with either Scramble (SCR) or siPool and stimulated or not with TGF-β1 and proteome profiling of FLS culture supernatant was performed. Among the 21 cytokines with significative differences identified between all conditions, three distinct response patterns were observed (Fig. 8B and S5B). In panel 1, TGF-β1 effects (red SCR + TGF-β1) on 8 cytokines secretions (downregulated: DKK1, PTX3 and VCAM-1; upregulated: LIF, MCP-1, PDGF-AA, Thrombospondin-1, and VEGF) were maintained in HS3ST-silenced FLS (blue siPOOL+TGF β1), suggesting that these responses were controlled by TGF-β1 independently of *HS3ST* expression. In contrast in Panel 2, HS3ST silencing effects (green siPool) on modulation of CXCL10, CCL5 and MIC-1 secretion were maintained with TGF-β induction (blue siPOOL+TGF β1), suggesting a specific regulation of their secretion by HS3ST expression and independently to TGF-β1. Finally in panel 3, IL-6, IL-8 and IL-11 displayed a distinct response pattern: HS3ST silencing amplified TGF-β1 induced IL-6 and IL-8 secretion and prevented IL-11 induction, suggesting that secretion of these cytokines could be specifically regulated by TGF-β1 and modulated by 3S-HS expression. These results were confirmed by an individual Elisa dosage (supp Fig. S5D).

**FIGURE 8:**
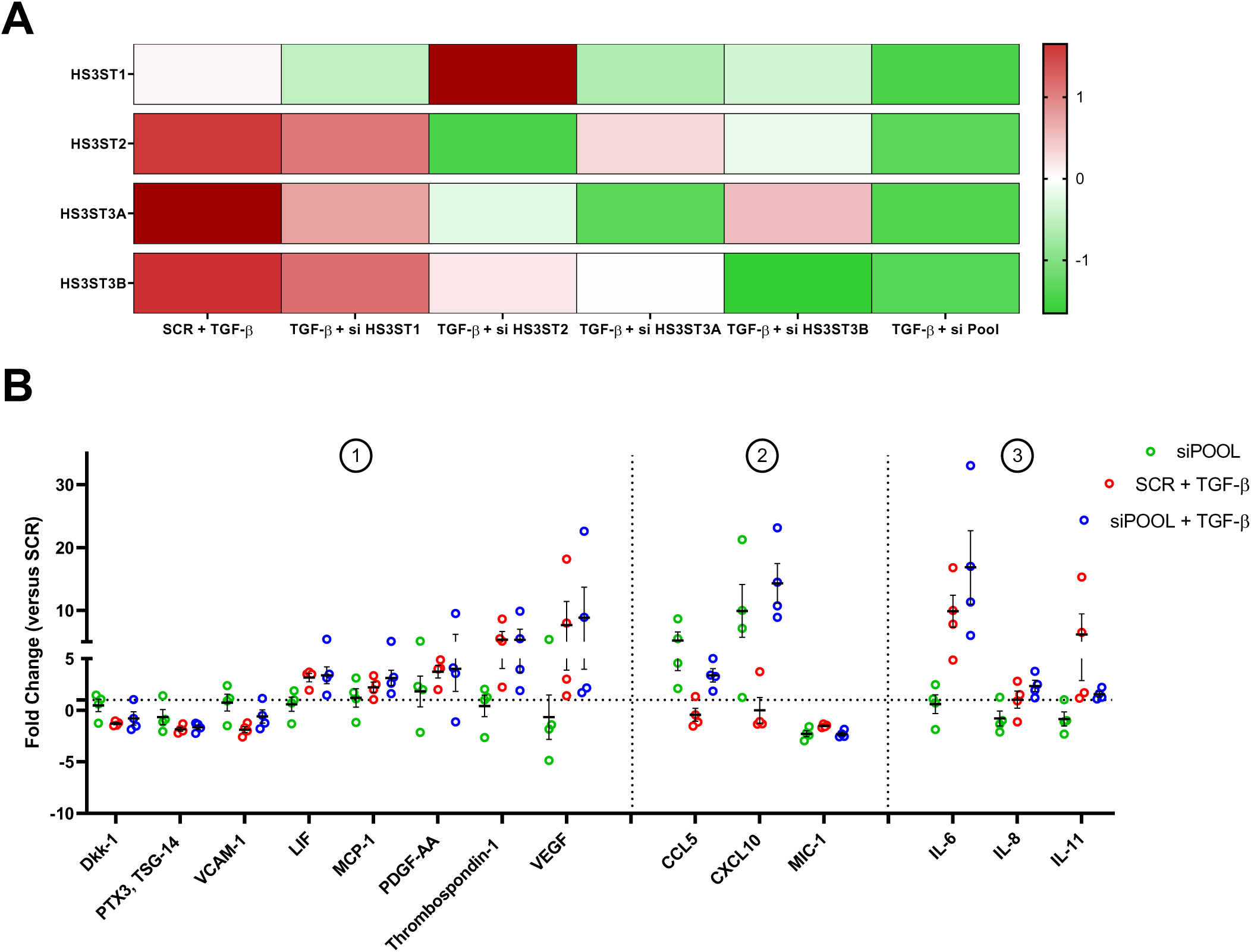
HS3ST knockdown and its impact on the TGF-β–induced inflammatory response in FLS. FLS were transfected and treated respectively with: scrambled siRNA (SCR) as control or 4 siRNA against HS3ST1, HS3ST2, HS3ST3, HS3ST3B isoforms individually or pooled (siPOOL) with or without TGF-β1 (5 ng·mL⁻¹). (A) Z-score heatmap showing relative mRNA expression of the 4 isoforms of HS3ST across 6 experimental conditions (z-scores computed per gene). Red indicates expression above the gene’s mean; green indicates expression below the gene mean. mRNA levels were normalized to α-tubulin and are shown as 2^−ΔCt^ values prior to z-scoring (see Methods). (B) Proteome array analysis of inflammatory mediators in FLS supernatants in SCR and siPOOL condition with or without TGF-β. Cytokines were grouped according to their response pattern: Panel 1, inflammatory mediators regulated by TGF-β independently of HS3ST silencing (DKK1, LIF, MCP-1, PDGF-AA, PTX3, thrombospondin-1, VCAM-1 and VEGF); Panel 2, inflammatory mediators regulated by HS3ST silencing independently of TGF-β stimulation (CCL5, CXCL10 and MIC-1); Panel 3, inflammatory mediators whose response to TGF-β was modulated by HS3ST silencing (IL-6, IL-8 and IL-11). Values are expressed as fold change relative to the SCR condition. Statistical analyses were performed using paired Wilcoxon signed-rank tests.

## DISCUSSION

We demonstrated for the first time that SF are enriched in GAG with relevant functionalities on different HBP and cells from joint, and with very specific 3S-HS sulfation pattern modulated by TGF-β. Then we try to bring some cues on cellular origins of HS from SF and their involvement in OA associated inflammatory response mediated by TGF-β.

### Sulfated GAG from SF are functional and modulate joint cells properties

SF-GAG are functional since able to bind HBP such as VEGF and FGF2 and to potentiate subsequent HBP’s receptor binding and transduced proliferation signal to fibroblast and endothelial cells. FGF2 signaling is known to be dependent to N- and 2-O-sulfated HS to transactivate FGFR receptor (Maccarana et al. 1993; Rusnati et al. 1994), whereas VEGF signaling is mainly modulated by NS- and 6S-HS (Robinson et al. 2006; Ferreras et al. 2012), suggesting that SF-GAG could contains a mix of these kind of sulfated motifs of HS. SF-GAG were able to modulate metabolic markers on murine chondrocytes also, probably through their ability to interact with HBP secreted by chondrocytes themselves. Among these chondrocytes HBP, we demonstrated that SF-GAGs interact not only with FGFs and VEGFs family members, but with numerous other ones involved OA pathophysiology such as Pleiotrophin (PTN), PDGF-BB, IL-6 and TGF-β1.

Elevated levels of PTN were described in OA cartilage at an early stage of the degenerative processes and involved in PG synthesis by chondrocytes (Tapp et al. 1999; Pufe et al. 2003; Kaspiris et al. 2013). PTN is expressed also in SF (Fig S1), on FLS and endothelial cells from inflamed synovial membranes, participating to matrix remodeling and neovascularization during chronic inflammation (Fadda et al. 2018). These effects are consistent with pleiotropic regulatory activities of PTN, including the modulation of VEGF-mediated angiogenesis through its receptors RPTPβ/ζ and N-syndecan (Imai et al. 1998; Papadimitriou et al. 2022; Choleva et al. 2024). Although PDGF-BB levels were low in OA SF (fig S1), PDGFs and PDGFR are expressed in both synovial fibroblasts and macrophages from OA SM (Pohlers et al. 2006). More recently, protective roles of PDGF-BB were demonstrated in cartilage degradation and subchondral bone remodeling during OA, as promoting chondrocytes anabolism and regulating inflammation markers (Zhu et al. 2021; Cai et al. 2023). Interestingly, these protective effects were associated to inhibition of VEGF mediated neovascularization that impact phenotype and properties of all the cells from the joint (Wang et al. 2025). Finally, we showed that SF GAG associated with proinflammatory TGF-β and IL-1b modulate their effect on FLS proliferation and IL6 and IL8 secretion *in vitro*, confirming functionality of these SF-GAG and HBP interactions in the joint.

Then, glycanic counterpart of synovia are not only degradative fragments associated to injury processes of cartilage but could be functional regulators of joint cell type’s properties: SF-GAG and specifically their HS fraction, could be new partners in the interplay of inflammatory/angiogenic factors such as TGF-β, VEGF, PDGF-BB and PTN. Moreover, as inter-donor heterogeneities were observed on binding properties and cells effect of SF-GAG, we proposed that GAG chemical structure need further exploration of their specific sulfated signature.

### GAG from OA SF are a mix of GAG species coming from different tissular origins

GAG extracted from human OA SF samples were much more concentrated than GAG from serum but not correlated with KL score in our study. Previous data tried to correlated SF-GAG to joint pathology severity according to KL-grade-evaluation in human OA samples (Kulkarni et al. 2016), on Gout and RA human samples (Carroll 1989) and in horses with distinct joint pathologies (Palmer et al. 1995). However, altogether these data suggest that the total GAG level is dynamically changing in SF and may not be the easiest biomarkers of joint pathology severity.

All these studies were focused on enriched CS, KS and HA fractions, that were suggested to originate from degradation of the most superficial layers of cartilage where PG loss (e.g. aggrecan) is most evident (Ratcliffe et al. 1993; Lohmander et al. 1993). Few data were delivered on HS from human SF and our study is the first to characterize this low fraction of HS species in total GAG extract from SF. Whereas fractions of GAG families from SF were similar to those of OA cartilage, our HPLC analysis showed very distinct sulfated pattern specifically for HS between OA SF and OA cartilage suggesting that they are distinct ones. In parallel, data strengthened the relevance of HS in synovial compartment. As example, conditional mouse embryo mutants lacking Ext1, an enzyme involved in HS synthesis, (created by mating Ext1f/f and Gdf5-Cre mice) display articulating superficial zone abnormalities with very low lubricin levels, indicating that local HS production is required to maintain the phenotype and function of joint-forming cells and (Mundy et al. 2011). Then, we hypothesize that HS from SF are not coming from cartilage degradation but could be produced by synoviocytes and confirmed this hypothesis by characterization of 3-*O* sulfated HS on FLS.

### HS from OA SF and FLS possess specific and rare sulfation pattern

HS from OA SF were mainly no sulfated chains, and their sulfated fraction was characterized by a very high level of tri-sulfated form of HS. Un-sulfated SF-HS fractions could be due to de-sulfated HS fragment from cartilage, obtain by matrix enzymes activities, such as Sulf endosulfatases (that remove 6-*O*-sulfate from HS) and heparanase (endoglycosidase that cleaves HS chains at low sulfation sites, releasing HS fragment that still retain biological activity (Vlodavsky et al. 2012)). Interestingly, Sulf are upregulated during OA and modulate BMP, FGF-2 and HS-Wnt-Frizzled signaling involved in joint (Otsuki et al. 2008, 2010). Moreover, their inactivation increases OA phenotype whereas Sulf intra-articular injections prevent synovial hyperplasia (Otsuki 2017). These suggest that regulation of 6-*O*-sulfated HS level in cartilage and SF, that serve as docking sites for numerous HBP (Lindahl et Li 2009), could be a key regulatory element of joint homeostasis and inflammation.

Inversely, the tri-sulfation pattern of SF HS, not observed on HS from cartilage, suggests that SF HS are not originate from cartilage degradation products or chondrocytes but could have another joint origin such as SM and FLS. Tri-sulfated HS signature could correspond to N-, 2*-O* and 6*-O* sulfated HS, or too much rarely associated with 3-*O* HS contents. We clearly identify this HS-3S sulfation signature on membrane of SF-derived and SM-derived FLS, associated to the heterogeneous expression of 4 *HS3ST* isoforms (*HS3ST2 < HS3ST1 < HS3ST3A* and *< HS3ST3B*). Then we confirmed that specific 3-*O*-sulfated HS in SF-GAG are coming from 3-*O*-sulfated HS of SF and SM derived Fibroblastic Like Synoviocytes. This is in accordance with the literature indicating that level and pattern of expression of *HS3ST* isoforms are respectively low and rare, restricted to endothelial cells from macro-vasculature, neuronal and glial cells, but also fibroblastic and immunogenic cells (Tiwari et al. 2006; Thacker et al. 2014). Moreover, associated patterns of 3-*O-*sulfated HS, that we clearly observed in OA FLS, remain always difficult to characterize according to their rarity and technical tools limits. Our data identify for the first time this rare glycanic signature on OA synovial compartment, and more specifically in FSL cells with immune phenotype. This suggests that 3-*O-*sulfated HS could be new biomarkers or actors of OA severity and/or synovitis, with biological roles and partners as well as associated inducible pathways that remain to be identified.

### TGF-β Drives a Robust Increase in 3S-HS Signal on FLS Membranes

Literature suggests that low basal levels of expressions of *HS3ST* and associated 3-*O*-sulfated HS are mainly induced in very specific pathological conditions (Denys et Allain 2019) and vulnerability (Ouidja et al. 2024). To explore signaling pathway involved in the joint synovitis, we performed sequence analysis of promoter region of *HS3ST* genes encoding all isoforms and identified at least consensus sequence for fixation of Smad transcription factors associated to TGF-β signaling (unpublished data). We focused on TGF-β signaling, according to its relevance during OA pathophysiology, and its experimental higher level in SF-OA and higher binding affinity to SF-GAG. Effectively, our data clearly demonstrated that TGF-β induced increase of 3-*O*-sulfated HS on FLS, by modulating the expression of 4 isoforms of *HS3ST* through the Smad 2/3 signaling pathway. Interestingly, TGF-β was already described as up-regulating the expression of *HS3ST3B1* in non-small cell lung cancer cells, while *HS3ST3B1* silencing reversed their mesenchymal phenotype induced by TGF-β (Zhang et al. 2018). More recently, TGF-β was involved in *HS3ST* expression in primary neurons from RTG4510 mice (Castillo-Negrete et al. 2025). It is also important to note that an increase in the expression of *HS3ST3B* was observed in many cell types exposed to inflammatory stimuli (Martinez et al. 2015; Sikora et al. 2016).

TGF-β1 is a central regulator in OA that displays both pro- and anti-inflammatory effects: while elevated levels promote synovial inflammation, hyperplasia, and fibrosis, it can exert anti-inflammatory actions also (Fava et al. 1991; Wahl et al. 1993; Diebold et al. 1995). Such complex effects could be explained by doses of active TGF-β available, and timing also that will transactivate different signaling pathways. In our study, we demonstrate for the first time that in FLS, TGF-β1 does not effectively signal through the Smad1/5/9 pathway, since phosphorylated SMAD1/5 remain sequestered in the perinuclear space, whereas clear activation and nuclear translocation of SMAD2/3 is involved. Moreover, TGF-β1 effects on *HS3ST* expressions on OA FLS were observed in a temporal manner: 4-hour of TGF-β1 treatment induced *HS3ST* expression while a 24-hour long treatment has an inhibitory effect (data not shown). Such temporal effect was already observed on chondrocytes *in-vitro,* since short TGF-β1 treatment induced PG synthesis whereas a treatment over several days induced MMPs the expression (Cherifi et al. 2021). Fibrosis, a hallmark of joint stiffness in OA, is also strongly driven by TGF-β in a time- and dose-dependent manner (Remst et al. 2014), whereas its inhibition prevents this process (Scharstuhl et al. 2003). As numerous matricial and pericellular HSPG are associated to the regulation of matrix sequestration of latent-TGF-β1, biodisponibility of its active form and activation of its receptors, its will be of great interest to establish a link between 3-*O*-sulfated HS and specific PG in this TGF-β induced context.

### 3-O-sulfated HS of FLS are involved in the control of their inflammatory secretome

Our data support the role of SF GAG and more specifically HS fraction, as functional regulators of joint cells properties. More precisely, we demonstrated that 3-*O*-sulfated HS modulates not only TGF-β-induced secretome from FLS, such as IL-6, IL-8 and IL-11 secretion, but reduced also the secretion of other relevant mediators during OA, such as CXCL10 and CCL5, independently of TGF-β. CXCL10 level are elevated in SF of OA patients (fig1S) and correlated with ‘activated’ phenotype of SF-derived cell (Trajerova et al. 2022), OA severity in mice model (Benigni et al. 2017) and immune infiltration (Yang et al. 2025). But CXCL10 role in OA SF is controversial since it’s a potent chemoattractant of subchondral mesenchymal progenitors, guiding them to the defect site and supporting repair with an effect on pain (Hernigou et al. 2021). CCL5 (RANTES) is highly expressed in OA SF (fig 1S) and FLS, and its interaction with its receptor CCR5 activates synoviocytes production of inflammatory mediators, triggering the immune response (C. Tang et al. 2010; Luo et al. 2024). CCL5 is a ligand of CCR1 also, influencing the polarization of M1 macrophage that will secrete CCL5 and CXCL10, thus promoting leucocyte infiltration and local inflammatory microenvironment in OA joint. Blocking CCL5-CCR1 axis reduce synovitis and alleviate OA (Xu et al. 2025) demonstrating that macrophages played a crucial role in the onset, progression, and recovery of OA within the synovium and joint space (Townsend 2014; Zhou et al. 2019). Then, 3-*O-*sulfated HS involvement in CXCL10 and CCL5 secretion by FLS in the SF could participate in the most global role of these chemokines on macrophage polarization and associated inflammation.

## Supporting information

Supp Datas

## Notes

### Competing Interest Statement

The authors have declared no competing interest.

### Funding

- Société Française de Rhumatologie
- Fondation Arthritis

