## Supplementary material for "Synovial fluid glycosaminoglycans signature involved 3-O-sulfated heparan sulfate modulated by TGF-β pathway in OA": Supp Datas

*** Co-authors**

Key words: Heparan Sulfate, 3-*O*-sulfation, Synoviocytes, TGFβ, Inflammation, Osteoarthritis,

Thanks to Société Française de Rhumatologie and Fondation Arthritis Grants

### SUPPLEMENTARY MATERIAL AND METHODS

**GAGs extraction and quantification from synovial fluid, serum and cartilage.**

Centrifuged synovial fluids (2,500 × g, 10 min, 4 °C) and non-centrifuged serum samples were mixed with a proteinase K–DNase buffer (2× for synovial fluid and 10× for serum) for subsequent GAG extraction, according to a modified version of the (Huynh, Villares, et al. 2012) protocol. Briefly, samples were successively treated with proteinase K (100 µg/mL, 56 °C 24h, Sigma-Aldrich) and DNase I (Qiagen;5 U/mL, 37 °C, overnight). Following digestion, samples were centrifuged (13,800 × g, 10 min, 4 °C) then filtered and mixed 1:1 with 4 M NaCl for 30 minutes. Proteins were precipitated with trichloroacetic acid (TCA, 10% Sigma-Aldrich) at 4 °C and lipids were removed from the resulting supernatant by two sequential chloroform extractions (1:1, v/v). The aqueous phase was dialyzed (Spectrapor) first against Tris buffer (50mM Tris-HCl, 50mM $CH_3COONA$, 2mM $CaCl_2$, pH 8), then against ultrapure water and frozen at −80 °C and lyophilized. Dried samples were reconstituted in ultrapure water to the initial sample volume (synovial fluid or serum) and stored at −20 °C until use. GAG were extracted form cartilage as previously described (Shamdani et al. 2020). Quality control (QC) samples were included in each extraction batch with pharmaceutic CS (Sigma). GAG were further used for quantification, HPLC analysis, Elisa competitive assay and functional tests.

**Dimethylmethylene blue (DMMB) assay of total sulfated GAGs**

Total sulfated GAGs extracted were quantified according to 1-9 dimethyl-methylene blue (DMMB, Sigma Aldrich) assay as described (Barbosa et al. 2003) according to a calibration curve of standard CS pharmaceutic (Sigma). For each sample, CS and HS amount were determined after specific digestion: CS were digested by Chondroitinase ABC (Sigma, 30 mU, 37°C , 1 hour), remaining HS and other GAG species were quantified by DMMB protocol and CS amount was calculated as the difference with total GAG quantified in the same sample. HS were chemically digested by nitrous acid treatment ($H_2O$. $NaNO_2$, 0.25 M, SIGMA S-2252) followed by addition of 33% acetic acid (VWR 20103.364) for 1 hour at RT and the reaction was stopped by 14% ammonium sulfamate (SIGMA 228745). Remaining GAG were quantified by DMMB protocol and HS amount was calculated as the difference with total GAG quantified in the same sample.

**Heparin/Glycosaminoglycan ELISA competition assay towards HBP**

The capacities of extracted GAG to interact with HBP were evaluated by an ELISA-based competition assay as described (Shamdani et al. 2020). The tested Recombinant Human HBP were VEGF-165 FGF-2 PDGFBB IL6 TGFβ and PTN (Supp Table S1), incubated with serial concentration of extracted GAG (from 0.01 to 100 µg/mL in PBS), or serial concentration of Heparin (Sigma) as a positive control, for 1h in ELISA 96-well plates previously coated with heparin-BSA complex. After washing steps, HBP remaining bound to heparin-BSA were targeted by specific antibody for 1h at RT, followed by a HRP-conjugated secondary antibody (Supp Table S1). Peroxidase activity was measured by TMB kit

(SI254783, Thermo Scientific, France). IC50 was defined as GAG concentration able to inhibit 50% of the HBP binding to immobilized heparin. As soluble Heparin was used as a positive control, its IC50 value was considered 100% of binding affinity. $IC_{50}$ of SF-GAG were normalized to heparin and inversely converted as % of Binding affinity to HBP (% of binding = 100 x ($IC_{50}$ -heparin / $IC_{50}$-HBP)).

**Human Umbilical Vein Endothelial Cells (HUVEC) Mitotic tests**

HUVECs were routinely maintained at 37 °C, 5% CO2 in full EBM medium (Lonza) according to provider protocol. For the mitogenic test, 5,000 HUVEC cells/well were seeded into 96-well plates in starving medium EMB-2 medium (Lonza) containing 0.5% FBS, without heparin and HBP overnight (37 °C, 5% CO2). GAG (5 ng/ml) were then added, in presence or absence of VEGF165 (5 ng/ml, Thermofisher RF233613) and incubated for an additional 72h. Then 25 µl of a 5 mg/ml MTT solution (Sigma M2128-10G) were added to each well and the plate was incubated for 3 hours (37°C, 5% CO2). Thereafter medium were taken out and 100 µL/well of DMSO:Sorensen's buffer solution (9:1, v/v, pH: 10.5,) was added, vortex to dissolve, absorbance determined at 570 nm (van Tonder et al. 2015). Heparin (5 ng/ml) was used as a positive control and considered as 100% effect.

**Baf32 cell mitogenic assay.**

BAF32 non adherent fibroblastic cells are deficient in HS biosynthesis and then need heparinoid compound provided in culture to facilitate formation of the FGF2-FGFR1-HS tertiary complex and transactivate FGF2 proliferative signaling. BAf32 cells were cultured in RPMI 1640 Glutamax medium (Gibco 61870-010), 10% SVF, Antibiotic-Antimycotic (Thermo 15240062) and 1 ng/mL final IL-3 (R&D systems 403-ML-010). Prior to mitogenic assay, cells were washed by centrifugation and resuspended in starving medium (RPMI 1640 Glutamax, 10% horse serum (Gibco 26050-070), antibiotic-antimycotic). Cells were seeded at 10.000 cells/well into 96 well plates in starving medium overnight (37˚C, 5% CO2). Thereafter extracted GAG and/or FGF2 (Invitrogen 68-8785-82) were added and incubated 72 hours (37˚C, 5% CO2). Cell viability was measured with a direct fluorometric detection (544 ex /590 nm) of the reduced products from PrestoBlue® (Invitrogen A13262), after 4 hours of incubation (37˚C, 5% CO2)

**HS and CS structure characterization by HPLC analysis**

The global HS and CS disaccharide composition were determined by HPLC, on total GAG samples treated with 20 mU of Chondroitinase ABC (SIGMA, 1h 30, 37°C) followed by heparitinases I, II and III (Iduron HEP-ENZ I, II, III) cocktail treatment (0.25mU of each, overnight at 37°C). In parallel, on the same sample, CS was specifically analyzed by digesting total GAG samples with Chase ABC and without treatment with the heparitinases cocktail. This allowed determination of the HS disaccharide composition by subtraction of the CS peaks from HS+CS signals and was performed to avoid chromatography column clogging resulting from CS digestion that interfered with HS analysis. Briefly, after filtration, samples were applied to a SAX-NP5 column on a HPLC system set as previously described (Huynh, Morin, et al. 2012): samples were automatically loaded onto the column and eluted by a solvent gradient consisting of eluent A (NaCl 20 mM) and eluent B (NaCl 1 M) at 1 ml/min flow rate. The gradient was 0–5 min (100% A), 5-25 min (from 100% to 0% A by steps), 25–29 min (0% A), 30–37 min (100% A). The column effluent was mixed (1:1) in line with 2% 2-cyanoacetamide and 25 mM NaOH, at 0.5 mL/min each. The mixture passed through a reaction coil (10 m length, 0.5 mm internal diameter) set in an oven at 120 °C, followed by a cooling coil (3 m length, 0.25-mm internal diameter). The effluent was monitored at 346 nm of excitation and 410 nm of emission with a fluorescence detector (JASCO; FP-1520). Peaks were assigned using commercial HS and CS disaccharides standards (Iduron). Moreover, 3 controls of digestion using commercial GAG were included in each experiment to validate complete enzymatic digestion. Results were expressed as % of area of each peak to the sum area of all peaks.

**Phenotypic characterization of FLS cultures**

The purity of the cultured SF-FLS and SM-FLS populations was assessed for all passages (0 to 7) and for every patient prior to experimental use by morphology (optical microscopy), adhesion to tissue culture plastic, and cell-surface marker profiling by flow cytometry. For immunophenotyping, SF-FLS and SM-FLS were enzymatically dissociated, washed and resuspended in fluorescence-activated cell sorter scan (FACS) buffer (PBS 1X, 1% BSA). FLS (60,000 – 100,000 cells) were incubated with fluorochrome-conjugated antibodies against FLS markers (CD55-FITC, CD44-PE, CD90-PE) and macrophage markers (CD14-FITC, CD68-Alexa647) at 1:100 dilution for 30min at room temperature in the dark. After three washes with FACS buffer (300 x g, 5min) cells were resuspended in 200µl PBS. A minimum of 50,000 singlet events were acquired per sample. Results are reported as the percentage of marker-positive viable cells. Data were analysed with FACS MACSQuant Analyser (Miltenyl Biotec).

**FLS proliferation assay**

FLS between passages 2 and 4 were seeded in 96-well plates at a density of 2,000 cells per well, corresponding to the optimal cell density previously determined using proliferation standard curves. Cells were cultured in complete medium for 24 h and subsequently synchronized for 24 h in serum-starved medium (DMEM low glucose with 1% PS). FLS were then stimulated in serum-starved medium with increasing concentrations of TGF-β (1, 5 and 20 ng/mL), FGF2 (1, 5 and 25 ng/mL), IL-1β (0.1, 1 and 10 ng/mL) or FGF18 (1, 10 and 100 ng/mL). Following identification of the concentrations inducing the strongest proliferative response, cells were stimulated with the selected growth factor or cytokine in the presence of increasing concentrations of OA synovial fluid-derived sulfated GAG extracts (0.01, 0.1 and 1 µg/mL). After 48 h of treatment, cell proliferation was assessed using the PrestoBlue™ Cell Viability Reagent (FisherScientific) according to the manufacturer's instructions.

Briefly, 10 µL of reagent was added to each well and signal intensity was measured after 4 h of incubation using a TECAN microplate reader at 570 nm with a reference wavelength of 600 nm. Results were normalized to the untreated control condition and expressed as fold change of control proliferation.

**Murine chondrocytes phenotypic analysis**

Primary murine articular chondrocytes were isolated from 6-day old mice as described by (Gosset et al. 2008). Chondrocytes were cultured for 6 days and then stimulated for 24h with OA-SF GAG (2.5µg/mL) or recombinant mouse interleukin-1β (IL-1β, R&D systems, 1ng/mL). Total RNA was extracted using Qiazol reagent (QIAGEN) and reverse transcription was performed using the high-capacity cDNA reverse transcription Kit (Applied Biosystems). Real-time PCR was performed using SYBR Green Master Mix (Applied Biosystems) to analyze the pattern of mRNA expression of anabolic (ACAN), catabolic (MMP13) marker (Supp Table S2). Relative gene expression was calculated using the $2^{-\Delta Ct}$ method, with RPL13A used as the reference gene. Data were analysed using AriaMx Software (Agilent Technologies).

**Secreted profiles in SF and FLS-conditioned media**

Biochemical analysis of cytokines and growth factors concentrations in SF and supernatant of FLS cultures, after treatments with selected mediators and/or GAG and/or siRNA was performed using PTN ELISA kit (Invitrogen, EH370RB) and DuoSet ELISA kits (R&D Systems) for IL-1β (DY201), IL-6 (DY206), TNF-α (DY210), IL-8 (DY208), CCL17 (DY364), TGF-β (DY240), VEGF (DY293B), FGF2 (DY233), PDGF-BB (DY220), CXCL10 (DY266), CCL5 (DY278), IL11 (DY218) according to the manufacturer's instructions. Analysis of FLS secretory profile after siRNA treatment was performed using Proteome Profile Human Cytokine Array kit (R&D systems ARY022B) according the manufacturer's instructions. Images were captured using a ChemiDocTouch Imaging system (Bio-Rad Laboratories) and were analysed using the Protein Array Analyzer plugin for ImageJ (2010 version available online: https://image.bio.methods.free.fr/ImageJ/?Protein-Array-Analyzer-for-ImageJ.html&lang=en). Spots of interest were normalized to an internal positive control after subtraction of representative background sample.

**Western Blotting**

FLS were lysed in RIPA buffer (ThermoFischer Scientific) supplemented with protease and phosphatase inhibitor cocktail (Sigma). Protein concentration was determined by BCA assay (Invitrogen), and equal amount of protein (20µg) were mixed with Laemmli and reducing buffer, boiled 5min at 95°C and resolved on a 10% or 12% SDS-PAGE gels. Proteins were blotted to PVDF membrane (Biorad) by liquid transfer overnight at 4°C. Membranes were blocked in TBST buffer (5 % milk in 25 mM Tris–HCl, pH 7.4, and 0.1% Tween) 1 h RT, and subsequently incubated with primary antibodies, diluted in TBST 5% BSA, overnight at 4°C: phospho-Smad2/3 Smad2/3, phospho-Smad1/5/9, Smad1, and β-actin (Supp. Table S3). After washing with TBST, immunoblots were incubated with peroxidaseconjugated anti-Rabbit or anti-Mouse (Supp. Table S3) 1h at RT. After washing, immunoblots were detected using the Luminata Forte Western Blot Substrate (Sigma) and chemiluminescence was captured with a ChemiDoc Imaging System (Biorad). Band intensities were quantified with ImageJ software and the results were expressed as a ratio of phospho-Smad/total-Smad expression.

**Quantitative analysis of HS3STs gene expression pattern in FLS by RT-qPCR**

Total cellular RNA was extracted from FLS using the RNeasy Mini Kit (Qiagen) according to the manufacturer's instructions, including on-column genomic DNA digestion with RNase-free DNase (Qiagen). Reverse-transcription was performed using random hexamers and the SuperScript First-Strand Synthesis System (Invitrogen). PCR amplification of cDNA was performed on an AriaMx thermocycler using SYBR Green DNA polymerase (Agilent Technologies) under the following conditions: initial denaturation (3 minutes, 95 °C), followed by 40 cycles of denaturation (95°C, 5 seconds) and annealing/extension (60 °C, 10 seconds). Relative gene expressions of the HS3STs isoforms (Supp Table S1) were determined using the ΔΔCt method (Livak et Schmittgen 2001) with α-tubulin used as the reference gene. Data were analysed using AriaMx Software (Agilent Technologies).

**siRNA transfection on FLS**

FLS were transfected with Lipofectamine™ 3000 (Invitrogen™ L300015) transfection reagent, and 20nM per siRNA against HS3T isoforms (Dharmacon HS3ST1 gene ID 9957, HS3ST2 gene ID 9956, HS3ST3A gene ID9955, HS3ST3B gene ID 9953) according manufacturer's instructions. A non-targeting scrambled siRNA was included as control. Cells were maintained for 24h after transfection, then stimulated with 5ng/mL of TGF-β for 24h after. At the end of the TGF-β1 stimulation, culture medium was replaced with starvation medium (DMEM low glucose, 1% Penicillin Streptomycin, 0.1%BSA). Cell pellets were collected 4h after the medium change to assess knockdown efficiency by RT-qPCR analysis and culture supernatants were collected 24h after the medium change for secretome profiling by proteome array. All treatments were performed in biological replicates (n=3-4).

#### SUPPLEMENTARY TABLES AND FIGURES

##### Supplementary Table S1: Commercial references of recombinant human Heparin Binding Proteins and antibodies for Elisa competitive assay

| Recombinant HBP | Supplier | Catalog Number | Coating concentration (ng/well) | Primary antibody (Supplier) | Catalog Number | Working Dilution | Secondary antibody | Catalog Number | Working dilution |
|---|---|---|---|---|---|---|---|---|---|
| FGF2 | Preprotech | 100-18B | 1 | Mouse anti-FGF2 (R&D System) | MAB233 | 1/2000$^{e}$ | Donjey anti-mouse HRP (Jackson ImmunoResearch) | 715-035-151 | 1/5000$^{e}$ |
| Il-6 | Preprotech | 200-06 | 12 | Mouse anti-IL-6 (Preprotech) | 500-M06 | 1/500$^{e}$ | Donjey anti-mouse HRP (Jackson ImmunoResearch) | 715-035-151 | 1/5000$^{e}$ |
| TGF-β1 | Preprotech | 11-21C | 10 | Mouse anti-TGF-β1 (Preprotech) | 500-M66 | 1/250$^{e}$ | Donjey anti-mouse HRP (Jackson ImmunoResearch) | 715-035-151 | 1/5000$^{e}$ |
| VEGF-165 | Reliatech | 300-076 | 1.2 | Rabbit anti-VEGF (Preprotech) | 500-P010 | 1/2000$^{e}$ | Goat anti-rabbit HRP (Jackson ImmunoResearch) | 111-035-144 | 1/2000$^{e}$ |
| PTN | R&D System | 252-PL | 5 | Goat anti-PTN (R&D System) | AF-252-PB | 1/1000$^{e}$ | Rabbit anti-goat HRP (Jackson ImmunoResearch) | 305-035-045 | 1/1000$^{e}$ |
| PDGFBB | R&D System | 220-BB | 4 | Goat anti PDGFBB (R&D System) | AF200NA | 1/2000$^{e}$ | Rabbit anti-goat HRP (Jackson ImmunoResearch) | 305-035-045 | 1/1000$^{e}$ |

##### Supplementary Table S2: Primer sequences used for RT-qPCR analyses

| Species | Gene | Forward primer (5'→ 3') | Reverse primer ((5'→ 3') |
|---|---|---|---|
| Human | HS3ST1 | GAACGAGGTCCACTTCTTCG | TTCTCCACTGTGAGCTGGTG |
| Human | HS3ST2 | TCTCCAAGAAGCCCGACATC | CGCCACTGACGAAGTGAATC |
| Human | HS3ST3A | TCACAGGTGGGGTCCAAG | GTTCTCTCCGCCCGAGTT |
| Human | HS3ST3B | CATCATCGGACGTCATCTT | AAACACGTGCCAGGACAAAT |
| Human | HS3ST5 | CCTGCTTGAAATGCTGAACC | TACCACTCAATGCCCTTACC |
| Human | HS3ST6 | GAGCCCCACTTCAGGGACAGG | ATCTGCCCATCCAGGGTTCG |
| Human | TUBA1A | GCAACAACCTCTCCTCTTCG | GAATCATCTCCTCCCCCAAT |
| Mouse | ACAN | CAGGGTTCCCAGTGTTCAGT | CTGCTCCCAGTCTCAACTCC |
| Mouse | MMP13 | TGATGGCACTGCTGACATCAT | TGTAGCCTTTGGAACTGCTT |
| Mouse | RPL13A | GGATCCCTCCACCCTATGACA | AGCCGAACAACCTTGAGAGC |

#### Supplementary Table S3: Antibodies used for Cytometry, Western Blot and Immunofluorescent staining analysis.

| Primary antibodies | | | | | |
|---|---|---|---|---|---|
| **Target** | **Host species** | **Application** | **Dilution** | **Supplier** | **Catalog number** |
| 10E4 | Mouse | IF | 1/1000[e] | Amsbio | 370255-1 |
| EV3C3v | | IF | 1/200[e] | Kindly provided by Prof. Toin H. van Kuppevelt (Radboudumc, Nijmegen, The Netherlands) | |
| RB4EA12 | | IF | 1/200[e] | Kindly provided by Prof. Toin H. van Kuppevelt (Radboudumc, Nijmegen, The Netherlands) | |
| AO4BO8 | | IF | 1/200[e] | Kindly provided by Prof. Toin H. van Kuppevelt (Radboudumc, Nijmegen, The Netherlands) | |
| HS4C3 | | IF | 1/200[e] | Kindly provided by Prof. Toin H. van Kuppevelt (Radboudumc, Nijmegen, The Netherlands) | |
| Anti-VSV glycoprotein | Rabbit | IF | 1/5000[e] | Sigma-Aldrich Chimie | V4888 |
| SMAD1 | Rabbit | WB<br>IF | 1/1000[e]<br>1/150[e] | Cell Signaling | #6944 |
| phospho - SMAD 1/5/9 | Rabbit | WB | 1/750[e] | Cell Signaling | #13820 |
| SMAD2/3 | Rabbit | WB<br>IF | 1/1000[e]<br>1/150[e] | Cell Signaling | #5678 |
| phospho - SMAD 2/3 | Rabbit | WB | 1/750[e] | Cell Signaling | #8828 |
| β-actin | Mouse | WB | 1/5000[e] | Sigma-Aldrich Chimie | A5316 |
| CD44 PE | Mouse | FACS | 1/100[e] | BD Pharmingen | 550989 |
| CD55 FITC | Mouse | FACS | 1/100[e] | BD Pharmingen | 555693 |
| CD90 PE | Mouse | FACS | 1/100[e] | BD Pharmingen | 555596 |
| CD14 AlexaFluor 647 | Mouse | FACS | 1/100[e] | BD Pharmingen | 562690 |
| CD68 AlexaFluor 647 | Mouse | FACS | 1/100[e] | BD Pharmingen | 562111 |

| Secondary antibodies | | | | | | |
|---|---|---|---|---|---|---|
| **Target** | **Host species** | **Flurophore/HRP** | **Application** | **Dilution** | **Supplier** | **Catalog number** |
| Anti-mouse IgG | Donkey | AlexaFluor 488 | IF | 1/250[e] | FisherScientific | 10398212 |
| Anti-mouse IgG | Donkey | HRP | WB | 1/10000[e] | Jackson ImmunoResearch | 715-035-151 |
| Anti-rabbit IgG | Goat | HRP | WB | 1/10000[e] | Jackson ImmunoResearch | 111035144 |

### Supplement data, Fig. S1

## A

| Samples | Cartilage | | Serum | | Synovial Fluid | | Synovial membrane |
|---|---|---|---|---|---|---|---|
| | Control (n=6) | OA (n=11) | Control (n=21) | OA (n=12) | GAG extraction (n=52) | Cell Culture (n=55) | Cell culture (n=11) |
| **Age (years): median (Q1-Q3)** | 82 (77-90) | 73 (64-97) | 47 (23-69) | 62 (42-79) | 72 (30-91) | 59 (37-92) | 70 (48-96) |
| **Sex: Woman/all (%)** | 6/6 (100) | 5/11 (45.5) | 4/21 (19) | 5/12 (41.6) | 34/52 (65) | 33/55 (60) | 5/11 (45.5) |
| **BMI (kg/m²): mean ± SD (n)** | N.D | 29.2 ± 5.4 (5) | 26.7 ± 3.3 (7) | 29.9 ± 1.9 (5) | 27.4 ± 4.2 (29) | 27.3 ± 3 (14) | 28.4 ± (10) |
| **Duration of painful symptoms (years): mean ± SD (n)** | N.D | N.D | N.D | 5.6 ± (1.8) | 6.3 ± 3.5 (22) | | 5.2 ± 2.6 (5) |
| **Current treatment : yes/no/N.D** | N.D | 4/0/7 | N.D | 4/0/9 | 26/8/18 | 30/8/17 | 4/2/5 |
| *- Analgesics* | | 3/4 | | 0/4 | N.D | N.D | 0/4 |
| *- NSAIDs* | | 1/4 | | 0/4 | N.D | N.D | 0/4 |
| *- CTC injection* | | 1/4 | | 4/4 | 13/26 | 13/30 | 4/4 |
| *- HA infection* | | 1/4 | | 2/4 | 8/26 | 7/30 | 1/4 |
| *- PRP injection* | | 0/4 | | 0/4 | 10/26 | 11/30 | 0/4 |
| **Kellgren-Lawrence classificaiton: mean ± SD (n)** | N.D | 3.9 ± 0.3 (11) | N.D | 2.5 ± 1(6) | 3.2 ± 0.97 (34) | 3.2 ± 0.6 (38) | 3.4 ± 0.5 (5) |
| **Lesquesne score: mean ± SD (n)** | N.D | N.D | N.D | 10.8 ± 4.7 (10) | 12.4 ± 3.6 (32) | 10.8 ± 2.4 (31) | 9.8 ± (1.8 (4) |

## B

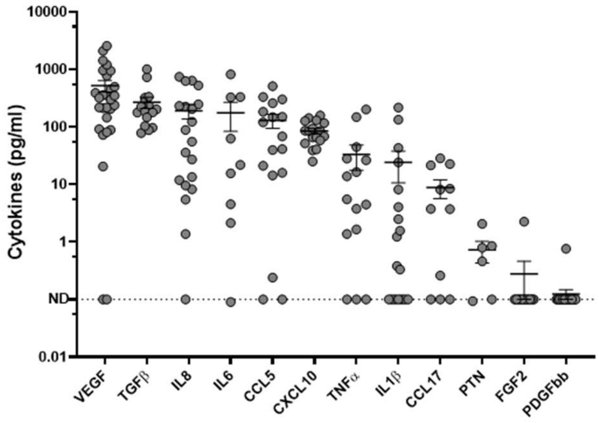


## C

| Correlation (GAG vs inflammatory mediators) | Rho (Spearman) | p-value | n |
|---|---|---|---|
| VEGF | -0,04511 | 0,8502 | 20 |
| TGF-β | -0,2377 | 0,3568 | 17 |
| IL-8 | 0,04167 | 0,876 | 17 |
| IL-6 | -0,1735 | 0,5193 | 16 |
| CCL5 | -0,1 | 0,7241 | 15 |
| CXCL10 | -0,1385 | 0,6375 | 14 |
| TNf-α | -0,1685 | 0,5475 | 15 |
| Il-1ß | 0,3497 | 0,22 | 14 |
| CCL17 | 0,2844 | 0,3966 | 11 |
| FGF2 | -0,2739 | 0,6667 | 9 |
| PDGFBB | -0,3039 | 0,2202 | 18 |

#### Supplement data, Fig. S2

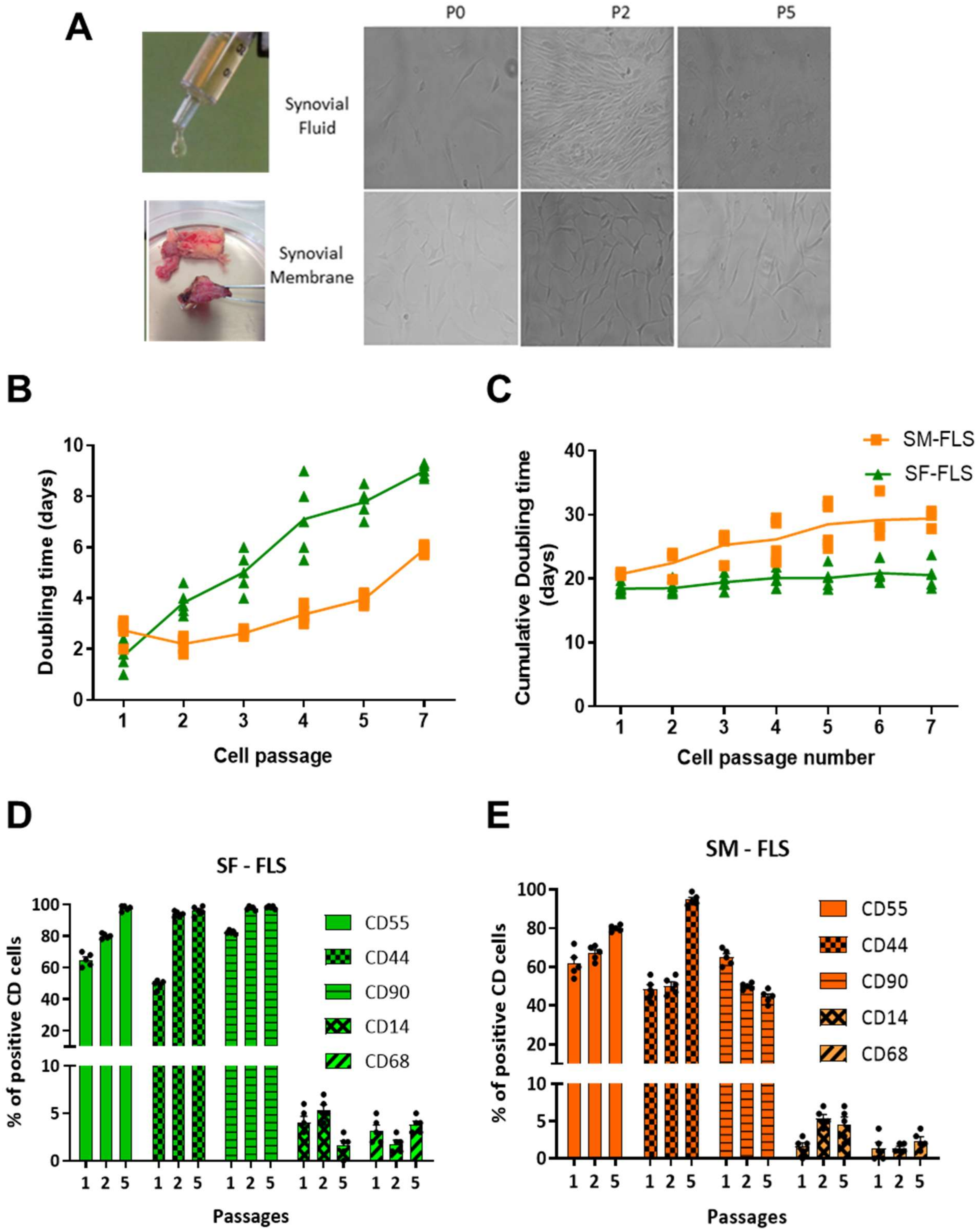

### Supplement data, Fig. S3

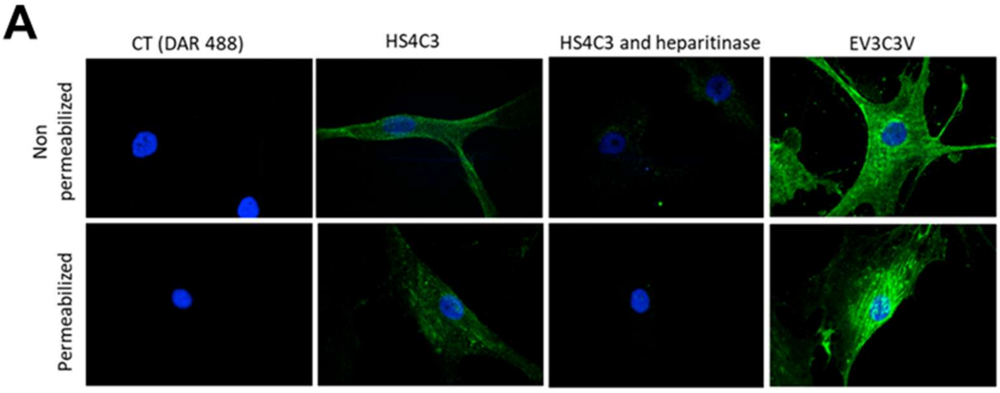

### Supplement data, Fig. S4

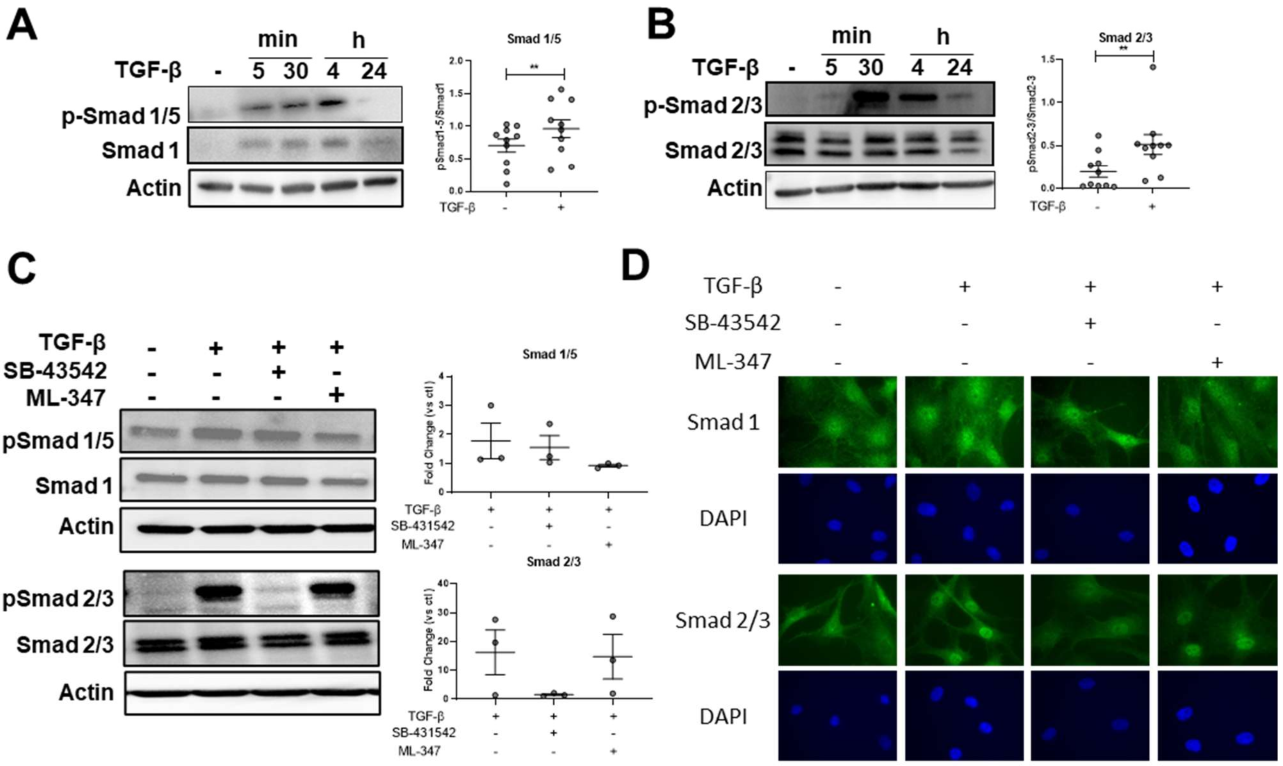

### Supplement data, Fig. S5

**A**

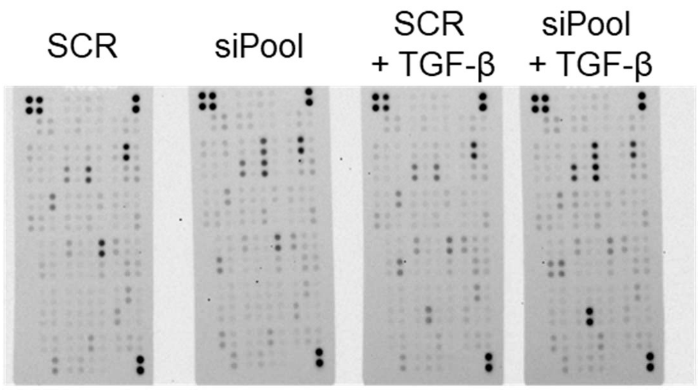


**B**

| Cytokine | SCR vs SCR TGF-β1 | SCR vs siPool | SCR vs siPool + TGF-β1 | SCR + TGF-β1 vs siPOOL | SCR + TGF-β1 vs siPool + TGF-β1 | siPool vs siPool + TGF-β1 |
|---|---|---|---|---|---|---|
| Angiogenin 283 | 0.282 | ***0.021*** | 0.282 | 0.114 | 0.686 | 0.343 |
| CCL5 | 0.282 | ***0.021*** | ***0.021*** | ***0.029*** | ***0.029*** | 0.343 |
| CHI3L1, YKL-40 | ***0.021*** | ***0.021*** | ***0.021*** | 0.343 | 0.561 | 1.000 |
| CXCL10 | 0.282 | ***0.021*** | ***0.021*** | 0.057 | ***0.029*** | 0.343 |
| Dkk-1 | ***0.021*** | 1.000 | 0.282 | 0.114 | 1.000 | 0.2 |
| Epidermal Growth Factor | 1.000 | ***0.021*** | 0.278 | ***0.029*** | 0.885 | ***0.029*** |
| FGF-19 | 0.282 | ***0.021*** | 0.282 | 0.245 | 0.343 | 0.486 |
| IGFBP-3 | ***0.021*** | ***0.021*** | ***0.021*** | 0.114 | 0.343 | 1.000 |
| IL-11 | ***0.021*** | 0.282 | ***0.021*** | ***0.029*** | 0.343 | ***0.029*** |
| IL-31 | 0.282 | ***0.021*** | 0.282 | 0.2 | 0.885 | 0.343 |
| IL-6 | ***0.021*** | 1.000 | ***0.021*** | ***0.029*** | 0.343 | ***0.029*** |
| IL-8 | 1.000 | 0.282 | ***0.021*** | 0.2 | 0.343 | 0.057 |
| LIF | ***0.021*** | 0.62 | ***0.021*** | ***0.029*** | 0.772 | 0.057 |
| MCP-1 | ***0.021*** | 0.282 | ***0.021*** | 0.486 | 0.486 | 0.2 |
| MIC-1 | ***0.021*** | ***0.021*** | ***0.021*** | 0.114 | ***0.029*** | 0.885 |
| PDGF-AA | ***0.021*** | 0.282 | 0.282 | 0.686 | 1.000 | 0.486 |
| PTX3, TSG-14 | ***0.021*** | 0.282 | ***0.021*** | 0.2 | 0.686 | 0.2 |
| Thrombospondin-1 | ***0.021*** | 0.62 | ***0.021*** | ***0.029*** | 1.000 | 0.057 |
| TNF-α | ***0.021*** | ***0.021*** | 0.282 | 0.343 | 0.343 | 0.057 |
| VCAM-1 | ***0.021*** | 1.000 | 0.282 | 0.057 | 0.2 | 0.343 |
| VEGF | ***0.021*** | 0.282 | ***0.021*** | 0.114 | 0.885 | 0.114 |

**C**

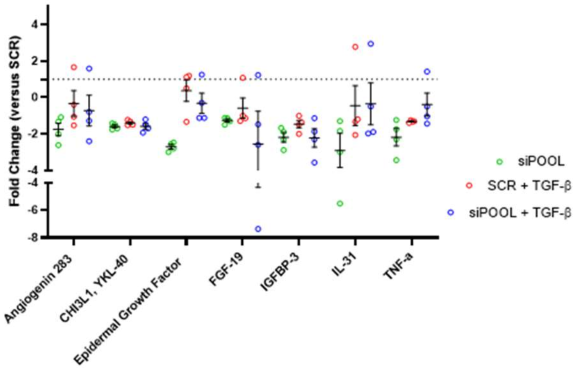


**D**

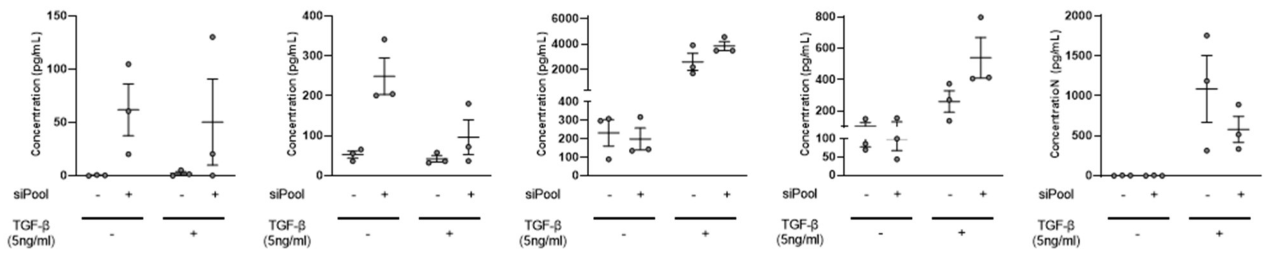

#### SUPPLEMENTARY FIGURES LEGENDS AND ASSOCIATED RESULTS

**FigS1: Patient characteristics and inflammatory profiles in synovial fluid from OA patients.** Patient characteristics. A total of 52 OA synovial fluid (SF) samples were included for GAG extraction and FLS cultures, whereas FLS culture from OA SM were established from 11 patients. 33 serum (CT n=21, OA n=12) and 17 cartilages (CT n= 6, OA n=11) samples were included also. Abbreviation: OA: osteoarthritis; GAG: glycosaminoglycan; BMI: body mass index; NSAIDs: non-steroidal anti-inflammatory drugs; HA: hyaluronic acid; PRP: platelet rich plasma; N.D: not determined. Median of age, sex distribution and BMI were similar between all these groups, without significant differences between CT and OA samples. The OA severity evaluated by Kellgren-Lawrence classification was moderate among serum and synovial fluid, whereas it was severe for cartilage and synovial membrane samples. B) Concentrations of secretory mediators in Synovial Fluid (SF) from OA patients. Results were obtained by ELISA dosage and shown as mean +/- SEM in pg/ml. All these mediators are heparin binding growth factors and cytokines (HBP), known to be involved in key inflammatory, proteolytic, angiogenic and reparative pathways during OA (De Roover et al. 2023; Boffa et al. 2021). VEGF, TGF-β, Il-8, Il-6, CCL5 and CXCL10 are among the highly concentrated, whereas CCL17, Il-1β, TNFα and PTN were lower. PDGFbb and FGF2 were not detected or only in some samples. In the case of interleukins IL-8, -6, -1β, VEGF and CCL5 and TNFα, 2 subgroups of levels were observed, suggesting that inflammatory profiles of patients could be different. (C) Relationship between SF-GAG and inflammatory mediator's concentrations were evaluated by Spearman's rank correlation (pvalue and n reported).

**Fig. S2: Phenotypic Characterization of Fibroblastic-Like Synoviocytes isolated from synovial fluid and synovial membrane.** (A) Representative phase-contrast images of primary FLS isolated from synovial fluid (SF) and synovial membrane (SM). Cell colonies were observed at passage 0 (P0, left panels), early passages (P2, middle panels), and later passages (P5, right panels). FLS isolated from SF or SM sources are spindle-shaped cells that progressively develop broader cytoplasmic processes, with morphological maturation occurred earlier in SF-derived FLS (passages 3–4) than in SM-derived FLS (passages 5–6). (B). Doubling time (DT) of FLS isolated from SF (n=5) and SM (n=5), calculated for each passage. (C) Cumulative population doubling time of SF- and SM-derived FLS (n=5). SM-FLS showed shorter population doubling times than SF-FLS, consistent with faster proliferation (D) and (E) Flow cytometry analysis of FLS isolated from SF (D) and SM (E) cultures (n=5). Expression of fibroblastic markers (CD90, CD44, and CD55) and macrophage-associated markers (CD14 and CD68) was assessed at passages 2–5. Cells consistently expressed FLS markers (CD55, CD44, CD90) up to passage 5 and lacked macrophage markers (CD14, CD68).

**Fig. S3: Validation of HS4C3 staining specificity on FLS.** Representative immunofluorescence images of FLS stained with the HS4C3 antibody recognizing 3-O-sulfated HS motifs, performed on non-permeabilized and permeabilized cells pretreated or not with a mix of Heparitinase I, II, and III prior to HS4C3 staining. Identical fluorescence intensity and distribution patterns indicate that HS4C3 epitopes are accessible at the cell surface. Enzymatic digestion of heparan sulfate chains completely abolished the HS4C3 fluorescent signal, confirming the specificity of the antibody for HS structures No differences in 3-O-sulfated and 2-O-sulfated HS motifs level or distribution were observed after permeabilization confirming the accumulation of these HS motifs on FLS surface (Fig. S3 A).

**Fig. S4: TGF-β selectively activated the SMAD2/3 pathway in FLS.** (A) and (B) Representative Western Blot of Smad 1/5 and Smad 2/3 expression and phosphorylation respectively, in the time course of FLS stimulated with TGF-β1 (5 ng·mL$^{-1}$) (Left panel) confirmed activation of the canonical TGF-β signaling pathway in FLS. Quantification (band intensities) of pSmad 1/5/9 : total Smad1/5 ratio and pSmad 2/3 : total Smad 2/3 ratio at the 30-min time point permits to identify this timing as one of the optimal. Data are expressed as mean ± SEM (n = 9) (right panel). (C) Representative western blots of SMAD1/5 and SMAD2/3 phosphorylation in FLS treated 30min with TGF-β1 (5 ng·mL$^{-1}$) in the absence or presence of SB431542 (10 μM) or ML-347 (1 μM) (left panel). Quantification of band intensities expressed as fold change versus untreated control (mean ± SEM, n = 3). Ratios were calculated as phospho-SMAD / total-SMAD and permitted to confirm specificity of activation. Paired t-test was used for Statistical analysis of western blot quantification. $p < 0.05$; $p < 0.01$.(D) Representative immunofluorescence images of SMAD1 (green) and SMAD2/3 (green) in FLS treated for 30 min with vehicle (control) or TGF-β1 (5 ng·mL$^{-1}$) ± SB431542 (10 μM) or ML-347 (1 μM). Nuclei are counterstained with DAPI (blue). In basal conditions, nuclear translocation of SMAD2/3 and SMAD1/5/9 were diffuse in cytoplasm with some perinuclear staining (Fig. S4D). Treatment with TGF-β1 induced a clear nuclear translocation of SMAD2/3 labeling, that was prevented with SMAD2/3 pathway inhibitor (SB431542) and not modified by the SMAD1/5/9 pathway inhibitor (ML347). By contrast, there was no modification of SMAD1/5/9 staining intensity or localization in presence of TGF-β and/or inhibitors, suggesting a selective activation of the SMAD2/3 pathway in FLS cells treated with TGF-β

**Fig. S5: Complementary analysis of cytokine secretion following HS3ST silencing and TGF-β1 stimulation in OA FLS.** (A) Representative images of Human Cytokine Proteome Profiler™ arrays (R&D Systems) performed on FLS supernatants treated with control (SCR) or HS3ST-targeting siRNA (siPOOL), with or without TGFβ stimulation (5 ng/mL, 24 h). Each membrane displays duplicate antibody spots for 105 cytokines. Representative results illustrate reproducible signal intensity patterns between patients (n=4). (B) Summary table of pairwise statistical comparisons (paired nonparametric tests, Wilcoxon signed-rank) between experimental conditions on higher expressed secretory mediators. (C) Proteome array analysis of inflammatory mediators in FLS supernatants following siRNA treatment (SCR or siPOOL) with or without TGF-β (5 ng·mL$^{-1}$). Inflammatory mediators shown in this figure exhibited significant variations between experimental conditions but were not included in the main categories described in Figure 8B because their regulation could not be unambiguously attributed to HS3ST-dependent, TGF-β-dependent, or cooperative TGF-β/HS3ST-dependent mechanisms. Values shown are fold-changes versus scrambled control (SCR). (D) Validation of selected targets (CCL5, CXCL10, IL-6, IL-8 and IL-11) by ELISA Dosage in FLS supernatants after siRNA treatment ± TGF-β (5 $ng \cdot mL^{-1}$). Data are shown as concentration in pg/ml (mean ± SEM).
